# Biases introduced by Ficoll-based isolation in acute myeloid leukemia sample analyses support the use of hemolysis

**DOI:** 10.64898/2026.02.11.705243

**Authors:** Bianca E Silva, Alison Daubry, Charline Faville, Adrien De Voeght, Jacques Foguenne, Mégane Jassin, Oswin Kwan, Leslie Correia Da Cruz, Gabriele Carriglio, Sébastien Charles, Frédéric Baron, Jo Caers, André Gothot, Grégory Ehx

## Abstract

Acute myeloid leukemia (AML) is a heterogeneous malignancy whose characterization relies on immunophenotyping and molecular profiling. While hemolysis is recommended for leukocyte isolation in clinical diagnostics, Ficoll-based density gradient centrifugation is widely used in research and biobanking. Here, we evaluated the impact of Ficoll isolation on commonly performed analyses of AML samples. Ficoll altered flow cytometry-based characterization by systematically enriching lymphocytes and AML blasts while depleting granulocytes. The increased T-cell content impaired AML engraftment in NSG mice, as T cells mediated terminal graft-versus-host disease. Although Ficoll had minimal impact on *ex vivo* AML blast expansion or chemotherapy response, RNA sequencing identified 1,136 differentially expressed genes compared with hemolysis, with Ficoll-processed samples notably leading to an overestimation of leukemic stem cell gene set expression. Immunogenomic deconvolution highlighted that Ficoll leads to an overestimation of CD8^+^ T-cell and monocyte abundances in sequenced samples. Mutation calling from RNA-seq data revealed substantial discrepancies between methods, including failure to detect a clinically relevant DNMT3A R882 mutation in a Ficoll-processed sample. Together, these findings support the systematic use of hemolysis to preserve cellular diversity and avoid unpredictable biases introduced by Ficoll-based isolation.

## INTRODUCTION

Acute myeloid leukemia (AML) encompasses multiple subtypes characterized by the clonal proliferation of myeloid precursors (blasts) exhibiting various degrees of differentiation^1^. Diagnosis and classification into AML subtypes and risk groups rely on the identification of multiple immunophenotypic and genetic markers, such as CD34 and CD117, mutations in FLT3-ITD, NPM1, and DNMT3A, as well as KMT2A (MLL) rearrangements^2^. Given the high heterogeneity of AML and the complexity of the diagnostic process, strict guidelines, established by the European LeukemiaNet (ELN), International Consensus Classification, and the World Health Organization, must be carefully followed to ensure diagnostic accuracy and consistency across clinical centers^1-3^.

Immunophenotyping by flow cytometry is a pivotal technique for characterizing AML and monitoring measurable residual disease (MRD)^4^. While analytical procedures on untouched blood or bone marrow samples are under development, standard analyses require the removal of red blood cells to enrich for leukocytes and enable antibody staining. The currently recommended method for this enrichment is hemolysis (recommendation B4 of the ELN2021^5^ and others^6, 7^), which involves selectively lysing red blood cells using a hypotonic solution while sparing leukocytes, which tolerate osmotic stress more efficiently. This technique therefore preserves the entire leukocyte population without altering the relative abundance of subpopulations, making it the gold standard for routine clinical flow cytometry. Density-gradient centrifugation represents the main alternative to hemolysis for leukocyte isolation. In this approach, blood is layered over a polysaccharide-rich solution (most commonly Ficoll-Paque); upon centrifugation, red blood cells and granulocytes sediment, while peripheral blood mononuclear cells (PBMCs) remain at the plasma–Ficoll interface and are collected. While hemolysis is widely used in clinical diagnostics for AML, Ficoll separation is more commonly employed in biobanking^8, 9^, large-scale multi-omics studies (e.g., Leucegene^10-13^ and BEAT-AML^14^), and fundamental research^15-17^.

The discrepancy in leukocyte isolation methods between clinical practice and experimental research prompted us to investigate the potential bias introduced by Ficoll-based isolation on the characterization of AML samples and downstream molecular analyses. In particular, the depletion of granulocytes by this method may lead to an overestimation of the abundance of smaller cell populations, such as T cells. In the era of immunogenomic profiling using bulk RNA sequencing (RNA-seq) to assess immune cell infiltration in bone marrow samples^18-20^, such bias could have unforeseen consequences. Additionally, although AML development in immunodeficient mice xenografted with human leukemic cells has traditionally been attributed to leukemic stem cells (LSCs)^16^, this concept has been challenged by experiments demonstrating leukemia-initiating activity outside the LSC fraction^21-24^. Therefore, to maximize the representativeness of the leukemia reconstructed in xenografted animals, it may be preferable to transplant non-fractionated leukemic samples.

In this study, we demonstrate that Ficoll isolation consistently enriches AML blasts and lymphocytes while depleting granulocytes. Although this effect was observed in every patient analyzed, its magnitude varied substantially from sample to sample. This introduced significant distortions in several downstream analyses, including engraftment in xenograft models, *ex vivo* blast proliferation assays, transcriptomic and immunogenomic analyses, and detection of mutations. Our study advocates for the systematic use of hemolysis and provides recommendations tailored to each specific analysis to be performed on the isolated sample.

## METHODS

### Patient samples

Peripheral blood samples (∼50 mL) from AML patients were obtained after written informed consent. Characteristics of the patients and samples are listed in Table 1.

**Table 1.**
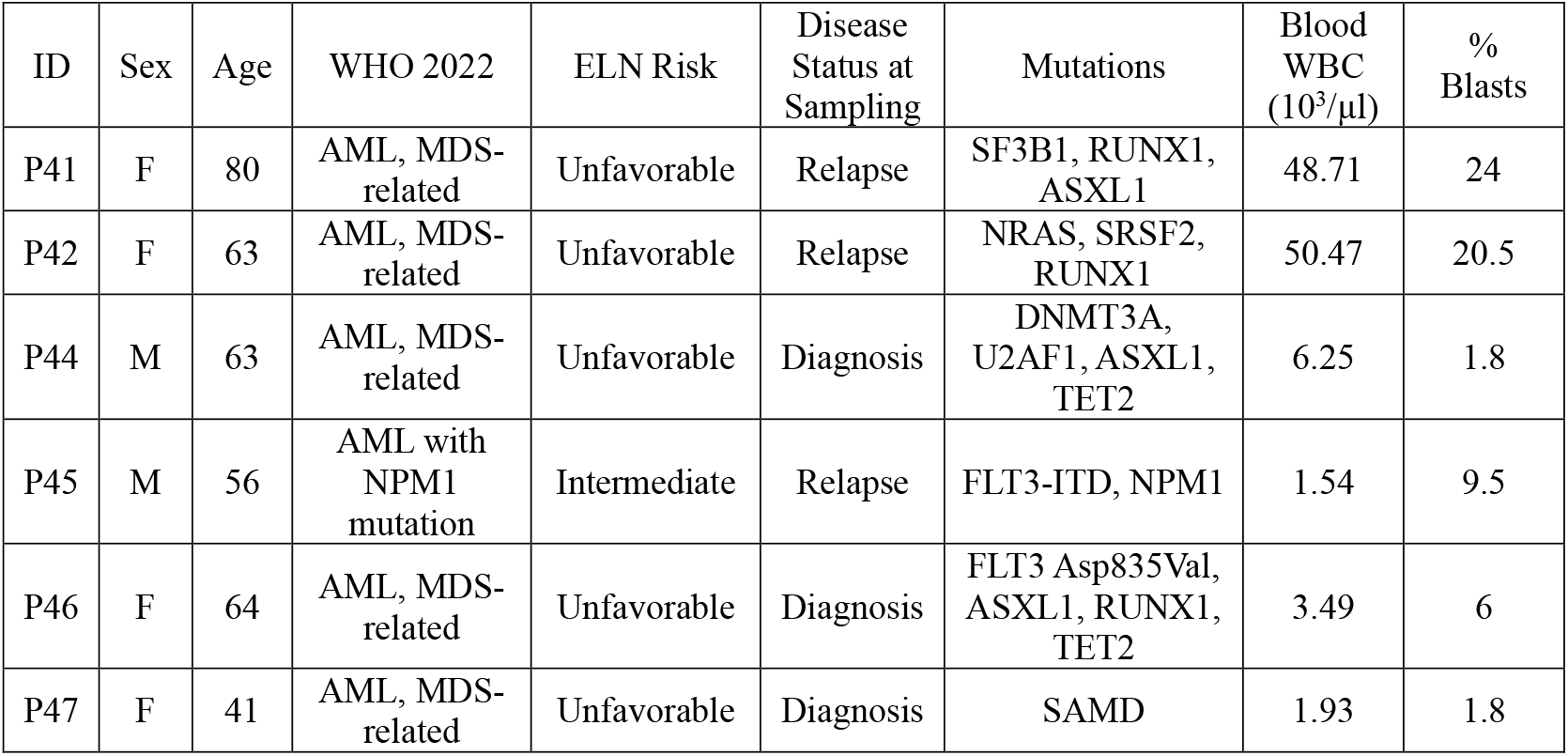
Characteristics of primary AML patient samples. White blood cell counts, blast percentages, and mutation data were obtained by our clinical department. Abbreviations: ELN, European LeukemiaNet; MDS, Myelodysplastic Syndrome; WBC, white blood cells; WHO, World Health Organization.

| ID | Sex | Age | WHO 2022 | ELN Risk | Disease Status at Sampling | Mutations | Blood WBC ($10^3/\mu\text{l}$) | % Blasts |
| --- | --- | --- | --- | --- | --- | --- | --- | --- |
| P41 | F | 80 | AML, MDS-related | Unfavorable | Relapse | SF3B1, RUNX1, ASXL1 | 48.71 | 24 |
| P42 | F | 63 | AML, MDS-related | Unfavorable | Relapse | NRAS, SRSF2, RUNX1 | 50.47 | 20.5 |
| P44 | M | 63 | AML, MDS-related | Unfavorable | Diagnosis | DNMT3A, U2AF1, ASXL1, TET2 | 6.25 | 1.8 |
| P45 | M | 56 | AML with NPM1 mutation | Intermediate | Relapse | FLT3-ITD, NPM1 | 1.54 | 9.5 |
| P46 | F | 64 | AML, MDS-related | Unfavorable | Diagnosis | FLT3 Asp835Val, ASXL1, RUNX1, TET2 | 3.49 | 6 |
| P47 | F | 41 | AML, MDS-related | Unfavorable | Diagnosis | SAMD | 1.93 | 1.8 |

### Leukocyte isolation

Equal volumes (∼25 mL) of each peripheral blood sample were processed either for red blood cell lysis (hemolysis) or Ficoll-Paque gradient centrifugation. Hemolysis was conducted by incubating one volume of blood with four volumes of lysis buffer^25^ (ammonium chloride (NH_4_Cl) 154.8 mM, EDTA 96.7 µM, potassium bicarbonate (KHCO_3_) 9.99 mM, diluted in ddH_2_O, pH 7.1, sterilized by autoclaving) for 15 min at room temperature under constant agitation. Leukocytes were then collected by centrifugation (500 ×g for 10 min) and washed in PBS. Ficoll-Paque gradient centrifugation was performed by layering blood samples (pre-diluted 2X in PBS) on 15 mL of Ficoll-Paque PLUS (cat: GE17-1440-02; Cytiva) and centrifuging for 20 min at 500×g without brake. PBMCs were collected by pipetting the mononuclear cell layer at the plasma–Ficoll interface and were washed with PBS. Isolated cells were counted with a Yumizen H500 hematology analyzer (Horiba).

### Immunophenotyping by flow cytometry

Cells were washed in PBS with 3% fetal bovine serum (FBS; cat: S-FBS-AU-015; Serana) before processing for flow cytometry staining. Cells were incubated with Fc receptor blocking solution (Human TruStain FcX, cat: 422302; Biolegend) for 10 minutes at room temperature before overnight incubation with surface antibodies at 4°C, as recommended in^26^. The following antibodies specific for human antigens were used (all purchased from Becton-Dickinson (BD) unless indicated otherwise): CD45-BV510 (clone HI30, cat: 563204); CD3-V450 (clone UCHT1, cat: 561812); CD19-APC (clone HIB19, cat: 17-0199-42; eBioscience); CD33-PE (clone HIM3-4, cat: 12-0339-42; eBioscience); CD15-FITC (clone W6D3, cat: 2215020; Sony Biotechnology); CD56-PE/Cyanine7 (clone HCD56, cat: 2191590; Sony Biotechnology); CD34-PerCP-Cy5.5 (clone 8G12, cat: 347222); CD117-APC-R700 (clone YB5.B8, cat: 565195). Cells were washed with PBS before analysis on a LSRFortessa flow cytometer (BD). Results were analyzed with FlowJo software v10 (Tree Star Inc.).

For mouse experiments, blood was collected by tail vein bleeding, lysed with RBC Lysis Buffer (cat: 00-4300-54; eBioscience), and washed in PBS with 3% FBS before staining. In addition to the above antibodies, the following antibodies were used: anti-human CD4-BV786 (clone SK3, cat: 563877); anti-human CD8-APC-Cy7 (clone SK1, cat: 348813); anti-mouse CD45-APC (clone 30-F11, cat: 559864). Cells were washed with PBS and incubated for 5 mins with 7-amino-actinomycin D (7-AAD, cat: 00-6993-50; eBioscience) before analysis on a cytoFLEX flow cytometer (Beckman Coulter (BC)).

Clinical routine analysis of immature myeloid populations was performed according to ELN recommendations^5, 27^ and included the use of the following antibodies specific for human antigens: CD34-FITC (clone 8G12, cat: 345801), CD13-PE (clone L138, cat: 347406), CD7-PerCPCy5.5 (clone M-T701, cat: 561602), CD33-PE/Cyanine7 (clone P67.6, cat: 333952), CD56-APC (clone NCAM16.2, cat: 341027), CD117-APC-A750 (clone 104D2D1, cat: B92450; BC), HLA-DR-Pacific Blue (clone Immu-357, cat: B36291; BC), and CD45-V500-C (clone 2D1, cat: 655873). Additional antigens were evaluated, such as anti-terminal deoxynucleotidyl transferase-FITC (HT1 + HT4 + HT8 + HT9, cat: IM3524; BC) and myeloperoxidase-PE (MPO; clone CLB-MPO-1, cat: B36288; BC), as well as CD300e-APC (clone UP-H2, cat: 656158), CD64-APCH7 (clone 10.1, cat: 561190), and CD36-V450 (clone CB38, cat: 561535) for determining monocytic lineage^2^.

### Mouse experiments

NOD-scid IL-2Rγ^null^ (NSG) mice (The Jackson Laboratory), aged 8–11 weeks, were administered an intravenous injection of 1×10^6^ leukocytes (isolated either by Ficoll or hemolysis) 24 h after 2.5 Gy total body irradiation. Graft-versus-Host Disease (GVHD) severity was assessed by a scoring system that incorporates 4 clinical parameters: weight loss, posture (hunching), mobility, and anemia, as previously reported^28^. Each parameter received a score of 0 (minimum) to 2 (maximum). Mice were monitored daily and scored for GVHD thrice weekly. Mice with a score of 6/8, weight loss >20%, or displaying signs of distress were sacrificed in agreement with the recommendation of the ethics committee. Final scores for dead animals reaching the ethical limit score were kept in the data set for the remaining time points (last value carried forward).

### *Ex vivo* culture of primary AML cells

Following isolation by Ficoll or hemolysis, primary AML cells were seeded at a density of 1×10^6^ cells/mL in an initial volume of 1 mL per well in 24-well plates in the presence of murine bone marrow stromal cells (OP9 cells; cat: CRL-2749; American Type Culture Collection) as described in^18^. Briefly, the OP9 feeder cells were pre-irradiated at 30 Gy and seeded at 80–90% confluence on 0.01% poly-D-lysine–coated plates (cat: A3890401; Gibco). Cells were transferred to a freshly irradiated OP9 feeder layer every 2 weeks or once the density exceeded 1.5×10^6^ cells/mL. Cultures were maintained in Iscove’s Modified Dulbecco’s Medium (cat: 12440053; Gibco) supplemented with 20% FBS, 1% penicillin-streptomycin (cat: 15140122; Gibco), and 50 µM of 2-mercaptoethanol (cat: 31350010; Gibco) at 37°C in a humidified incubator with 5% CO_2_. A cytokine cocktail, consisting of SCF (50 ng/mL, cat: 300-07), G-CSF (20 ng/mL, cat: 300-23), GM-CSF (20 ng/mL, cat: 300-03), FLT3-L (50 ng/mL, cat: 300-19), IL-3 (20 ng/mL, cat: 200-03), and IL-6 (20 ng/mL, cat: 200-06), was added twice weekly (all cytokines from Peprotech).

Cell proliferation and viability were monitored thrice week by 7-AAD exclusion on a cytoFLEX flow cytometer. Before each measurement, the wells were carefully flushed to recover as many blasts as possible without detaching the OP9 stromal layer. Cultures were reseeded to 5×10^5^ cells/mL once this density was exceeded. Each subculture was documented to calculate the cumulative cell count at the end of the experiment, using the formula: cumulative cell count = cell count on the day × cumulative number of previous subcultures performed. If the cell viability dropped below 60%, the cells were centrifuged at 500×g for 5 minutes and resuspended in fresh medium supplemented with the cytokine cocktail.

### Drug preparation

Cytarabine (AraC; cat: HY-13605 ; MedChemExpress) and cytokines were reconstituted in PBS to the desired stock concentration. Sterile solutions were aliquoted and stored at -80°C for up to six months.

### Analysis of senescence by flow cytometry

Cells were seeded at a density of 5×10^5^ cells/mL in 24-well plates and cultured in the presence of the cytokine cocktail but in the absence of the OP9 feeder layer. Immediately after seeding, the cells were treated daily with 1.5 µM AraC for three consecutive days. Staining with 5-Dodecanoylaminofluorescein Di-β-D-Galactopyranoside (C_12_-FDG; cat: HY-126839; MedChemExpress) was performed 24 h post-treatment with AraC. Briefly, the cells were washed by centrifugation (500×g, 5 min), and the pellet was resuspended in culture medium containing 100 nM bafilomycin A1 (cat: HY-100558; MedChemExpress), followed by incubation for 1 h at 37°C and 5% CO_2_. Subsequently, the cells were incubated for 2 h with 33 µM C_12_-FDG or DMSO (cat: 67-68-5; Sigma-Aldrich) for unstained control conditions. After two washing steps, 7-AAD was added immediately before flow cytometry acquisition to assess C_12_-FDG fluorescence (FITC channel) exclusively in viable cells.

### RNA sequencing

After isolation, one million leukocytes were cryopreserved at -80°C in TriPure Isolation Reagent (cat: 11667165001; Roche). Total RNA was isolated with the RNeasy Mini Kit (cat: 74106; Qiagen) according to the manufacturer’s instructions, and RNA quality and concentration were assessed using the 5200 Fragment Analyzer (Agilent Technologies). A total of 1 µg of RNA (RQN score >8) was processed using the NEBNext Ultra II Directional Poly(A) Library Prep Kit. Library profiles were verified on the Qiaxel Advanced (QIAgen) instrument using a DNA screening cartridge. The final dual-indexed libraries were quantified by qPCR using KAPA SYBR FAST (cat: KK4600 ; Roche) with Illumina standard and pooled equimolarly.

Sequencing was performed on the NovaSeq 6000 (Illumina) sequencer on an S4 flow cell, generating paired-end 150-cycle reads with a sequencing depth of 10 million fragments sequenced per sample. Raw sequencing data were demultiplexed using sample-specific indexes, filtered for quality, and converted into FASTQ files for downstream analysis using the bcl2fastq software. Sequencing data quality were assessed using the FastQC tool for individual reports per sample, and with MultiQC to generate a report comprising all samples.

### Bioinformatic analyses

Transcript expression quantifications were performed with kallisto v0.51.0^29^ in stranded mode. Kallisto’s transcript-level count estimates were converted into gene-level counts using the R package tximport v1.14.2. EdgeR v3.28.1 was used to normalize counts using the TMM algorithm and output counts per million (CPM) values. Differential gene expression analysis was conducted in R3.6.1 as reported previously^30^. In brief, raw read counts were converted to CPM, normalized relative to library size, and lowly expressed genes were filtered by retaining genes with CPM >0 in at least two samples using edgeR^31^ and limma^32^ v3.42.2. Subsequently, voom transformations and linear modeling using limma’s lmfit were performed. Moderated t-statistics were then computed with eBayes. To perform a paired differential gene expression analysis, a custom contrast matrix was built with the model.matrix function to compare each Ficoll sample to its Lysis equivalent per patient. The contrast matrix was then used as input to the contrasts.fit function of limma. Differential expression was defined using an adjusted p-value <0.05 and an absolute log_2_ fold-change (FC) ≥1.

Gene set enrichment analysis (GSEA) was performed with fgsea package v1.12.0 in R^33^. A pre-ranked gene list was generated by ranking expressed genes obtained from the paired limma–voom analysis on the moderated t-statistics. HALLMARK gene sets were obtained from the MSigDB database, and blood transcriptional modules (BTM) gene sets were obtained from^34^.

Immune cell population deconvolutions were performed with CIBERSORTx, the default LM22 matrix and 100 permutations. Mutation analyses were performed by aligning raw RNA-seq reads on the GRCh38.113 genome with STAR v2.7.11a^35^ running with default parameters to generate bam files. Single-base mutations with a minimum alternate count setting of 5 were identified using freeBayes v1.3.8^36^ and mutations were annotated with SnpEff v5.3a^37^ and the clinVar database (downloaded on 31-05-2025)^38^. Transcriptomic analyses performed on the BEAT-AML cohort^14^ were conducted on publicly available data (https://biodev.github.io/BeatAML2/). Raw counts were converted to CPM values with EdgeR as indicated above and used as input of CIBERSORTx.

For immunogenomics analyses, CIBERSORTx deconvolution outputs were aggregated to approximate the cell populations defined by flow-cytometric gating. CD8^+^ T cells were defined as SSC^low^CD3^+^CD8^+^, whereas CD4^+^ T cells (including CD4 memory, CD4 memory activated, and CD4 naive subsets) were defined as SSC^low^CD3^+^CD4^+^. B-cell fractions (naive B cells, memory B cells, and plasma cells) corresponded to SSC^low^CD19^+^ cells, while resting and activated NK-cell fractions corresponded to SSC^low^CD56^+^ cells. Monocytes were defined as SSC^int^CD14^+^ cells, and granulocytes (including neutrophils and eosinophils) were defined as SSC^high^CD15^+^ cells. Samples were subsequently ranked according to the abundance of each population (using flow-cytometry–derived proportions among CD45^+^ cells for Lysis samples, or aggregated CIBERSORTx values for Ficoll and Lysis samples), and rankings were compared between the two methods for each isolation approach.

### Statistics

Unless specified in the figure legends, paired t-tests were used to compare two conditions, and correlations were assessed with the Pearson correlation coefficient. Boxplots display the median with 25^th^ and 75^th^ percentiles and whiskers extending to the 10^th^ and 90^th^ percentiles. Bar plots show the mean ± standard deviation (SD), when indicated. Plots and statistical tests were mainly performed with GraphPad Prism v10.6. For all statistical tests, ^∗∗∗∗^ refers to p < 0.0001, ^∗∗∗^ refers to p < 0.001, ^∗∗^ refers to p < 0.01 and ^∗^ refers to p < 0.05.

## RESULTS

### Ficoll gradient isolation distorts leukocyte composition toward blasts and lymphocytes

To compare the impact of isolation methods on AML characterization, six peripheral blood samples were freshly collected from random AML patients. Patient characteristics are summarized in Table 1. Equal parts of each sample were processed using either hemolysis (referred to as the “Lysis” condition hereafter) or Ficoll-based isolation (“Ficoll” condition hereafter), and the resulting samples were analyzed by flow cytometry.

We first analyzed the effects of the isolation methods on leukocyte purity (CD45^+^ cells) and noticed that Ficoll samples were significantly enriched with leukocytes, while Lysis samples consisted of substantially more cell debris (Figure 1A). After gating on viable cells and excluding debris and doublets, we then analyzed the frequency of various markers within the leukocyte compartment (encompassing lymphocytes, monocytes, granulocytes, and AML blasts) and observed that Ficoll significantly increased the proportion of CD3^+^ T cells while depleting CD15^+^ neutrophils (Figure 1B). This imbalance resulted from the depletion of SSC^high^ cells (mainly neutrophils) and the selection of SSC^low^ cells (mainly lymphocytes and blasts) by the Ficoll method (Figure 1C). Consequently, the frequency of lymphocytes (SSC^low^CD45^high^) significantly increased after Ficoll isolation (Figure 1D). No significant impact was found on the frequencies of CD19^+^, CD56^+^, CD33^+^, and CD34^+^ cells (Figure 1B).

**Figure 1.**
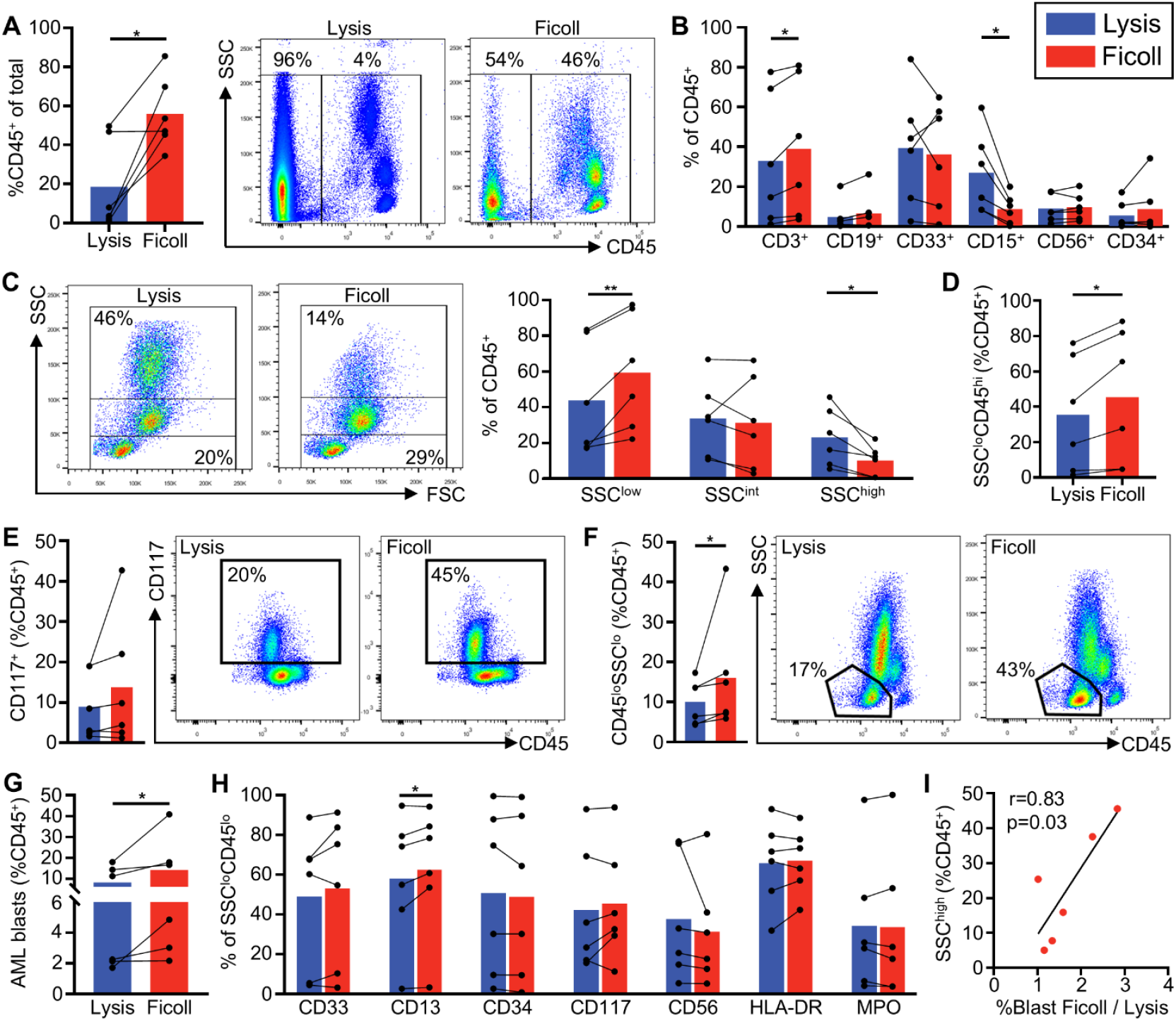
Selective recovery of AML blasts and lymphocytes with Ficoll isolation. Leukocytes were isolated from the peripheral blood of six AML patients by either hemolysis or Ficoll-based gradient centrifugation and analyzed by flow cytometry. Paired t-tests were used for comparisons, where each line represents a patient in the bar plots. **(A)** Frequency of leukocytes (CD45^+^ cells) among all detected events. **(B)** Frequency of the indicated markers among leukocytes. **(C)** Flow cytometry dot plot comparing the frequency of leukocytes segregated according to their granularity (side-scatter, SSC). **(D)** Frequency of lymphocytes (gated as SSC^low^CD45^high^ cells) among leukocytes. **(E)** Flow cytometry dot plot and comparison of the frequency of CD117^+^ cells among leukocytes. **(F)** Flow cytometry dot plot and comparison of the frequency of AML blasts (gated as SSC^low^CD45^low^ cells) among leukocytes. **(G)** Frequency of AML blasts computed through a Boolean clinical-grade gating strategy. **(H)** Frequency of the indicated markers among AML blasts (gated as SSC^low^CD45^low^ cells). **(I)** Pearson correlation between the frequency of cells with high granularity (SSC^high^, from panel C) in the Lysis condition and the enrichment of AML blasts (clinical frequencies from panel G; %blasts_Ficoll_ / %blasts_Lysis_) by the Ficoll approach. All bar plots show the mean (**p<0.01, *p<0.05; paired t-test).

In addition, Ficoll increased the frequency of CD34^+^ cells in four patients (Figure 1B). The analysis of CD117^+^ cell frequencies revealed a similar trend, with increases observed in the same four patients (Figure 1E). Because CD34 and CD117 are the most commonly used markers of AML blasts, we examined the frequency of these cells more specifically. We analyzed the frequency of SSC^low^CD45^low^ cells, an immature phenotype widely shared by AML blasts^39^, and observed that Ficoll significantly increased their frequency (Figure 1F). To validate this observation, we performed a clinical-grade analysis of the blasts based on a Boolean gating strategy using CD34, CD117, HLA-DR, CD13, CD7, CD56, and CD45 expression (ELN staining^1^). This confirmed the enrichment of AML blasts by the Ficoll method (Figure 1G). Given that blasts are SSC^low^ cells, this enrichment probably also results from the relative depletion of SSC^high^ cells by Ficoll. Importantly, no striking impact on the phenotype of blasts was highlighted by these analyses (except for a modest but significant effect on CD13), suggesting that the composition of the blast population is largely preserved in patients with conventional (SSC^low^) blasts (Figure 1H). However, substantial patient-specific changes in certain markers were observed, suggesting that Ficoll may alter blast phenotypes unpredictably in some cases. Interestingly, a significant correlation between the frequency of SSC^high^ cells in the Lysis samples and the degree of blast enrichment by the Ficoll method was observed (Figure 1I), suggesting that the Ficoll isolation method impacts the composition of the AML samples with abundant granulocytic cells.

Altogether, these results show that the isolation method significantly distorts the composition of AML samples through the depletion of granulocytes and the enrichment of lymphocytes and leukemic blasts.

### T-cell enrichment by Ficoll isolation impairs *in vivo* AML blast expansion

AML blast enrichment prior to transplantation is frequently performed to establish patient-derived AML xenografts in immunodeficient mice^15, 25, 40, 41^. However, the infusion of unfractionated Ficoll-isolated mononucleated cells is also performed^18, 42-46^. Given the differences observed between Lysis and Ficoll isolation methods on AML and non-AML cell frequencies, we reasoned that the isolation method may impact AML engraftment in immunodeficient mice. Using the cells of P46 for which we had enough material post-isolation, we transplanted one million leukocytes after Ficoll or Lysis isolation to three mice per condition (Figure 2A). In this patient, the CD45^+^ cell population presented similar frequencies of AML blasts (6.4% vs. 7.2% in Lysis and Ficoll, respectively) while Ficoll enriched for T cells by 15% (Figure 2B-C). Therefore, mice transplanted with AML cells isolated through Ficoll also received proportionally more T cells (4.5 ×10^5^) than Lysis-engrafted ones (3 ×10^5^). These T cell quantities are typically considered insufficient to induce xenogeneic GVHD upon injection to NSG mice^47^.

**Figure 2.**
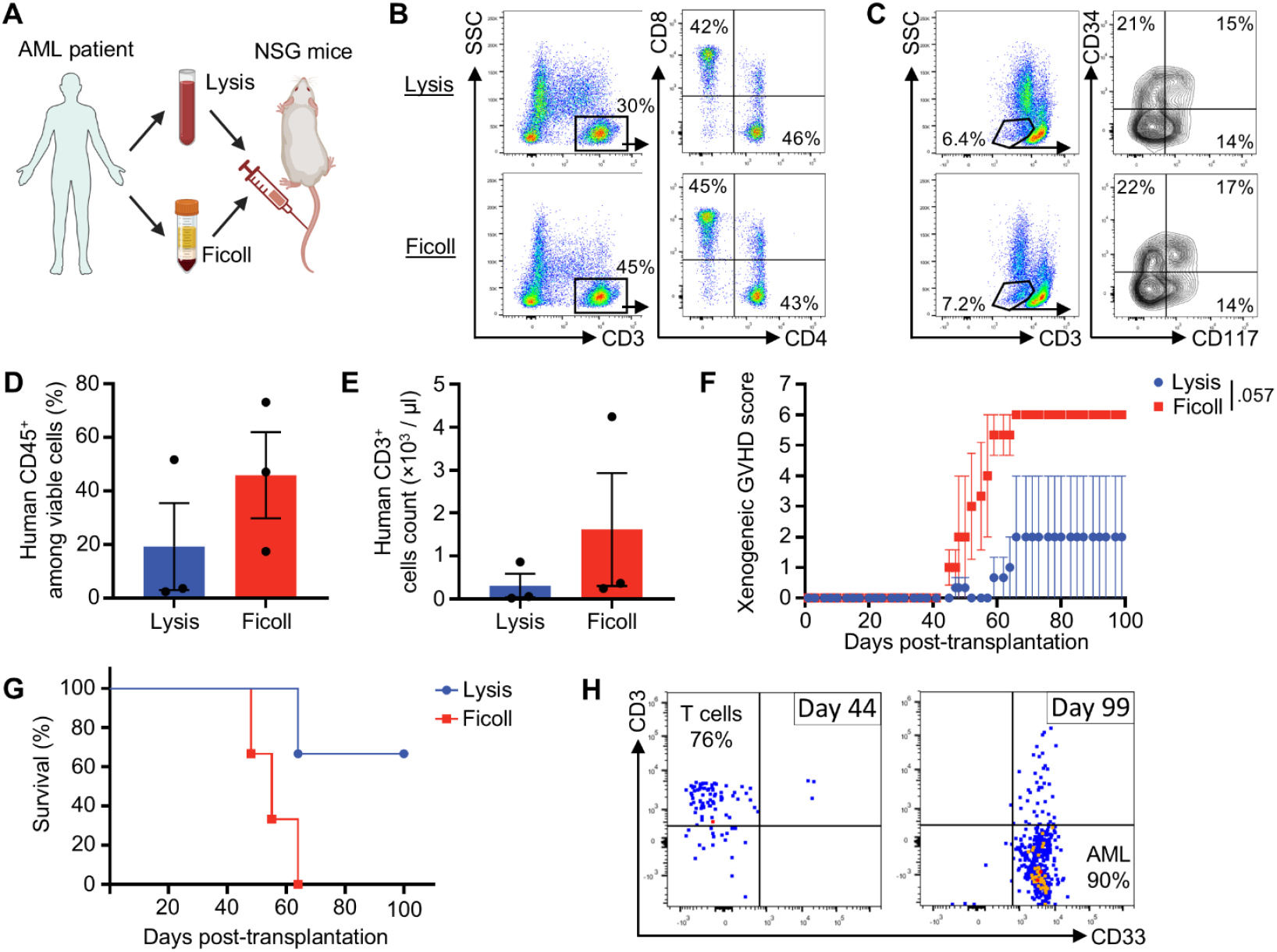
Impact of the isolation method on AML blast expansion *in vivo*. Leukocytes were isolated from the peripheral blood of one AML patient (P46) by either hemolysis or Ficoll-based gradient centrifugation. The total leukocyte fraction was injected into NSG mice 24 h after non-lethal irradiation. **(A)** Experimental design. **(B)** Flow cytometry analysis of the frequency of T cells (CD3^+^) and their subsets, CD4^+^ and CD8^+^ cells, among the leukocytes before injection into NSG mice. **(C)** Flow cytometry analysis of the frequency of AML blasts (SSC^low^CD45^low^) among leukocytes before injection into NSG mice, with CD34 and CD117 expression. **(D)** Frequency of human CD45^+^ cells among leukocytes (_human_CD45 + _mouse_CD45 cells) in the peripheral blood of NSG mice on day 42. **(E)** Concentration of human CD3^+^ cells in the blood of NSG mice on day 42. **(F)** Scoring curve of GVHD symptoms in NSG mice over the entirety of the experiment (ANOVA-2). **(G)** Log-rank survival curve; mice were sacrificed upon reaching an ethical score limit of 6. **(H)** Flow cytometry analysis of the frequency of T cells and myeloid cells in the blood of NSG mice on days 44 and 99.

Flow cytometry analyses showed that Ficoll mice presented a trend toward greater engraftment of human CD45^+^ cells and greater abundance of CD3^+^ T cells (Figure 2D-E). This higher T-cell abundance was likely responsible for the development of unexpected GVHD symptoms that we observed in the three mice transplanted with these cells (Figure 2F). These mice were euthanized between days 48-64 upon reaching the typical ethical limit score of GVHD symptoms (Figure 2G). In contrast, only one mouse transplanted with a Lysis sample required sacrifice on day 64 (due to GVHD), and the two other mice survived until the end of the experiment (day 100). Flow cytometry analysis on the peripheral blood of these animals showed that one of these two mice presented a high frequency (90%) of CD3^─^CD33^+^ cells, which were most likely AML cells since normal myeloid cells do not survive and expand in NSG mice^47^ (Figure 2H). This mouse presented only circulating T cells on day 44, suggesting that the failed long-term engraftment of T cells enabled the expansion of AML cells.

### Limited impact of Ficoll isolation on *in vitro* AML blast expansion

Cell lines do not reflect the epigenetic state of primary cancer cells^48^ or their mutation spectrum^49^. Therefore, primary cell culture of AML blasts is increasingly used, notably to study AML response to chemotherapy^42^. In light of our results, we assessed whether the isolation method could also affect the efficacy of such cultures. From the cells of four patients for which enough material was available, we performed *ex vivo* expansion of the total leukocyte population by culturing AML blasts on a layer of irradiated bone marrow stromal cells (OP9) in the presence of a cytokine cocktail (Figure 3A). The cells from two patients (P42 and P44) eventually proliferated, in agreement with the success rate (∼55%) of this culture system reported previously^17^.

**Figure 3.**
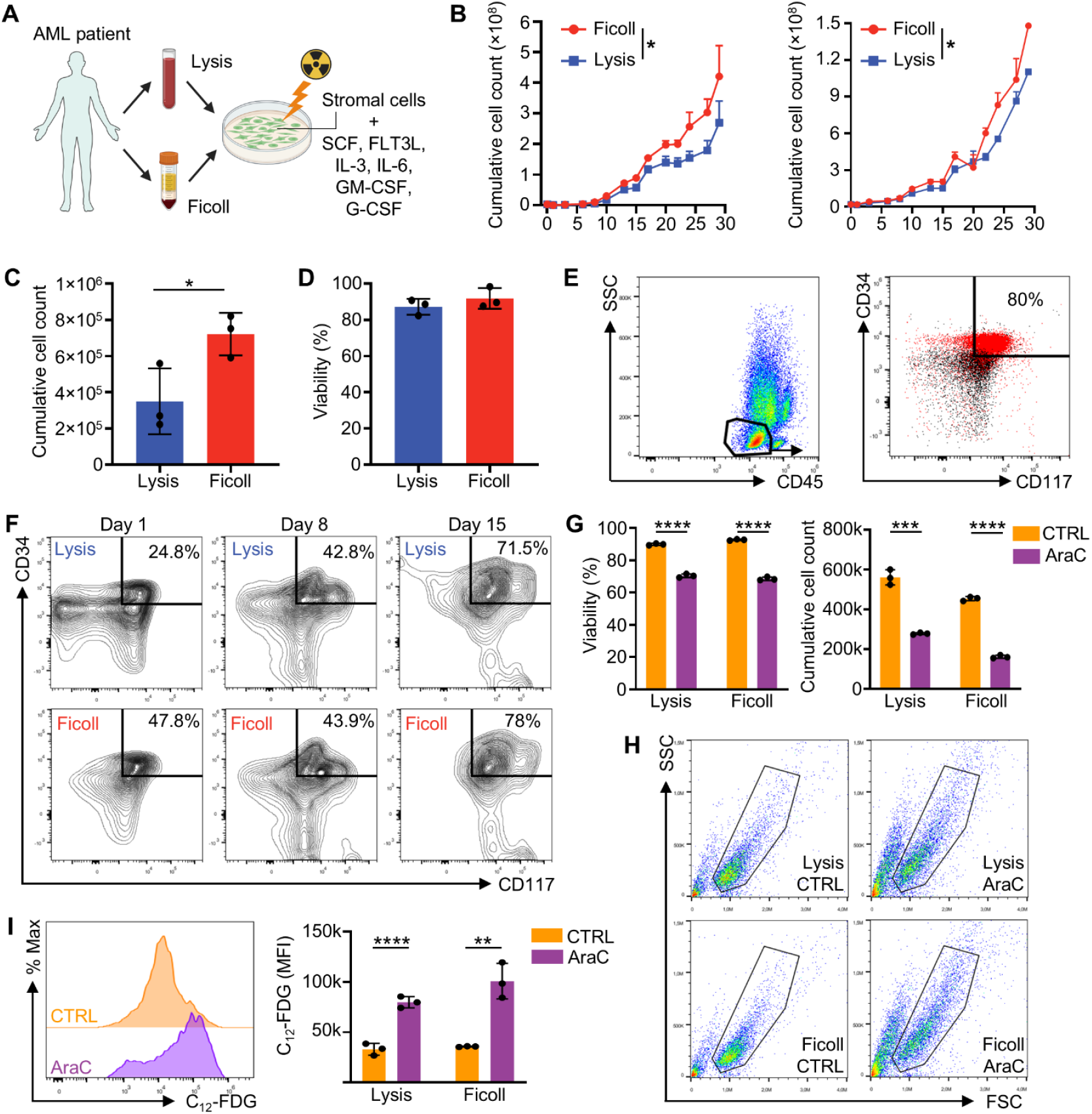
Impact of the isolation method on AML blast expansion *ex vivo*. Leukocytes were isolated from the peripheral blood of four AML patients by either hemolysis or Ficoll-based gradient centrifugation. AML blasts were then expanded *ex vivo* for four weeks. **(A)** Experimental overview. **(B)** Cumulative count of viable (7-AAD^─^) cells from patient 42 (left) and 44 (right) over the course of the culture. ANOVA-2 tests were used to compare expansion between the two conditions. **(C)** Cumulative count of viable (7-AAD^─^) AML blasts (SSC^low^CD45^low^) after 72 h of culture. **(D)** Viability (% of 7-AAD^─^) of AML blasts (SSC^low^CD45^low^) after 72 h of culture. **(E)** CD34 and CD117 expression of SSC^low^CD45^low^ blasts (in red) and non-blasts (in black) after 24 h of culture. The percentage represents the frequency of CD34^+^CD117^+^ cells among SSC^low^CD45^low^. **(F)** Evolution of CD34^+^CD117^+^ cells among CD45^+^ cells at indicated time points. **(G)** Viability (% of 7-AAD^─^) and cumulative cell count of cultured AML blasts treated *ex vivo* with 1.5 µM AraC for 72 h. **(H)** Cell size (FSC) and granularity (SSC) of AML blasts after 72 h of AraC treatment. **(I)** Flow cytometry histogram depicting SA-β-Gal activity using the fluorogenic β-galactosidase substrate C_12_-FDG, gated on viable cells (7-AAD^─^), after 72 h of AraC treatment. All bar plots show the mean ± SD (****p<0.0001, ***p<0.001, **p<0.01, *p<0.05; t-test).

Surprisingly, Ficoll-isolated AML cells proliferated slightly but significantly better than Lysis-isolated cells from both patients (Figure 3B). Since both patients displayed comparable results, we focused on patient 42, where the largest differences were observed (although all subsequent findings were also confirmed in patient 44). Flow cytometry analysis 72 h after the start of the culture showed that Lysis had lower SSC^low^CD45^low^ AML blasts (Figure 3C), in agreement with the lower starting percentage of blasts in these samples. No difference in cell death was observed between the two conditions (Figure 3D). Therefore, the slower growth in the Lysis condition probably resulted from the lower initial abundance of AML blasts rather than from a greater cell death by the hypotonic shock of the Lysis procedure.

We then questioned whether the isolation method might affect the immature/stem cell phenotype of the blasts over time and observed that the SSC^low^CD45^low^ blasts from this patient were CD34^+^CD117^+^ (Figure 3E). We therefore analyzed the frequency of CD34^+^CD117^+^ cells among total leukocytes (CD45^+^ cells) at three time points (days 1, 8, and 15) and observed that their frequency tended to increase with time (especially in the Lysis condition), showing that this population was expanding and that our culture system preserved their leukemic phenotype (Figure 3F). On days 8 and 15, no difference in blast frequency was found between Ficoll- and Lysis-isolated blasts. Altogether, these results suggest that the lower frequency of AML blasts in Lysis samples at the start of the culture is responsible for the mild delay of expansion, but this does not eventually impact the AML cell phenotype in the culture.

Next, we assessed whether blasts expanded from both isolation methods respond similarly to chemotherapy *in vitro*. AraC was tested since it is the main drug given as first-line chemotherapy to patients in the conventional 7+3 regimen. Blasts were collected from our primary cultures on day 21 of expansion and were treated with AraC. At a dose of 1.5 µM, AraC significantly reduced the cell counts and viability of AML cells (Figure 3G). In agreement with a previous report assessing AraC effects on primary AML cultures^42^, AraC increased the cell size and granularity (Figure 3H). Another key feature of the relapse-mediating AML post-chemotherapy is the expression of Senescence-Associated(SA)-β-Galactosidase^42^. We therefore stained our cells with the fluorogenic substrate of the SA-β-Gal, C_12_-FDG^50^, and observed that AraC increased its activity, as expected (Figure 3I). Importantly, no significant difference in AraC effect on SA-β-Gal was observed between isolation methods, in agreement with the comparable blast phenotype between conditions.

### The Ficoll isolation method alters the transcriptional profiling of AML samples

Over the last decade, multiple large-scale transcriptomic studies were conducted in AML by consortia such as TCGA, Leucegene, or BEAT-AML. Typically, such studies involve performing RNA-seq on unsorted leukocyte fractions. Frequently, the method to isolate these fractions is not specified, and the comparison of results from one cohort to another could be biased simply by the usage of divergent isolation methods. We therefore investigated whether the isolation method impacts the transcriptional profile of AML samples in bulk RNA-seq analyses.

RNA was extracted directly from the total leukocytic fraction isolated after the last wash from either the Lysis or Ficoll procedures, consistent with the approach used in the BEAT-AML program^14^. Samples from five patients had RNA quality scores sufficient to perform sequencing. Paired differential gene expression analysis showed that 1,136 genes were differentially regulated (p<0.05), with 152 being highly (FC > 2) enriched in Ficoll samples, and 59 in Lysis samples (Figure 4A and Table S1), revealing that the isolation procedure has a significant impact on the transcriptomic profile of AML samples. We then performed GSEA with HALLMARK^51^ and BTM^34^ gene sets. HALLMARK analysis highlighted higher normalized enrichment scores of inflammatory pathways in Lysis samples while proliferation-related pathways were enriched in Ficoll samples, likely reflecting the enrichment of AML blasts in these samples (Figure 4B). BTM analysis suggested an enrichment of neutrophils and inflammation pathways in Lysis samples, as well as an enrichment of erythrocytes, monocytes, and B cells with Ficoll (Figure 4C). The enrichment of HALLMARK inflammation pathways probably reflected the greater abundance of neutrophils in Lysis samples, as suggested by BTMs. The enrichment of B cells, monocytes, and erythrocytes in the Ficoll samples was in agreement with our flow cytometry results and the absence of a hemolysis step before RNA extraction in the Ficoll procedure.

**Figure 4.**
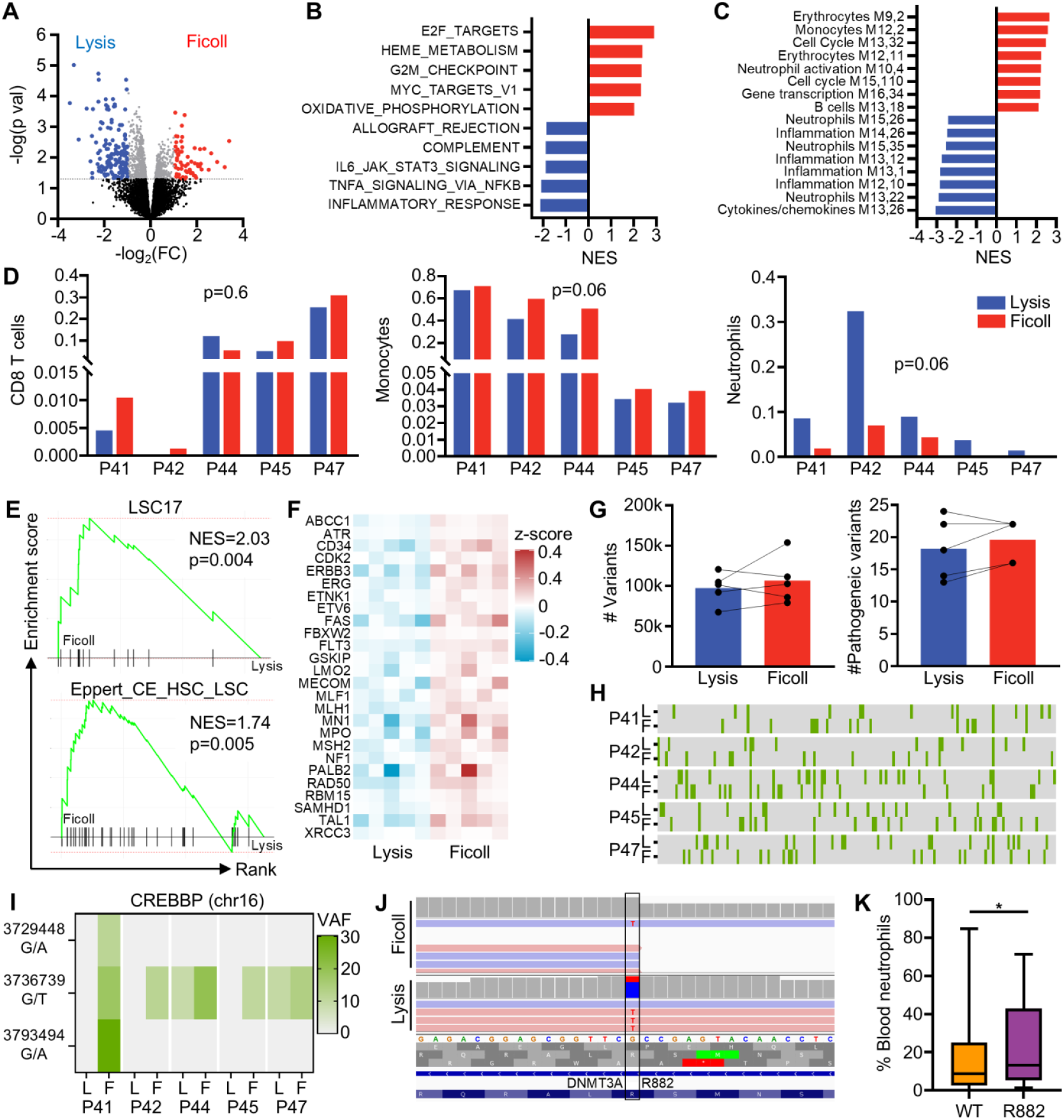
Transcriptional impact of the AML blast isolation method. Leukocytes were isolated from the peripheral blood of five AML patients by hemolysis or Ficoll-based gradient centrifugation. RNA-seq was performed on total leukocyte fractions. **(A)** Volcano plot of the paired limma-voom analysis performed on bulk RNA-seq data. Significant (p<0.05) genes are colored in red (log_2_(FC)>1), blue (log_2_(FC)<1) or grey. **(B, C)** Paired GSEA with the HALLMARK (B) or BTM (C) matrices. Gene sets upregulated in Ficoll samples are indicated in red and those upregulated in Lysis samples are indicated in blue. All gene sets are significantly (FDR<0.05) differentially regulated. **(D)** CIBERSORTx deconvolution analysis of the abundance of CD8^+^ T cells, monocytes, and neutrophils in RNA-seq data. **(E)** Paired GSEA of indicated gene sets. **(F)** Heatmap of z-scores of indicated genes in the Lysis and Ficoll samples. **(G)** Total (left) and pathogenic (right) variant counts in Ficoll and Lysis samples. **(H)** Heatmap of the distribution of pathogenic variants (each column is a different variant) in the five patients tested. L, Lysis; F, Ficoll. **(I)** Heatmap of the variant allele frequency (VAF) of the three mutations in the CREBBP gene in the five patients tested. L, Lysis; F, Ficoll. **(J)** IGV visualization of the R882 mutation in the DNMT3A gene of P44. **(K)** Flow cytometry analysis of neutrophils in the blood of DNMT3A R882-mutated patients in the BEAT-AML cohort (*p<0.05, Mann-Whitney test).

Because the two isolation methods yielded markedly different cellular compositions, we questioned whether these differences would be detectable in our RNA-seq data, as suggested by the BTM analysis. CIBERSORTx^52^ deconvolutions of the cell type composition indicated a greater abundance of neutrophils in Lysis samples and a trend toward enrichment for monocytes and CD8^+^ T cells with Ficoll (Figure 4D). Assuming that the enrichment of monocytes may result from the greater abundance of AML blasts, we performed a GSEA analysis with two gene sets specific to LSCs^53, 54^ and observed an upregulation of both gene sets (Figure 4E). Accordingly, the expression of many AML-related genes (such as CD34, FLT3, and MPO) was upregulated in Ficoll samples (Figure 4F). Altogether, these results show that the Ficoll isolation method leads to an overestimation of the AML gene expression, together with an underestimation of the abundance of normal immune cells.

### Ficoll isolation impacts the detection of mutations in RNA-seq samples

Analyses of mutation expression in cancer patient cohorts is frequently performed on bulk RNA-seq data. The expression of such mutations can then be related to patient survival^55^, neoantigen generation^56^, or the oncogenic process itself^57^. We therefore investigated whether the isolation method could have an impact on such analyses. After variant calling, a similar number of variants were detected between both methods (Figure 4G). An average of 32% of variants were commonly detected, and neither method enriched the number of specific variants. To discriminate the variants most likely to be relevant to AML biology, we annotated them with the ClinVar database^58^, and focused on pathogenic and likely pathogenic variants (Table S2 and Figure 4G). In agreement with an enrichment of blasts by the Ficoll approach, a trend toward more pathogenic variants was found in Ficoll samples. The identity of detected mutations changed dramatically between approaches, with an average of only ∼11% common between both approaches (Figure 4H), highlighting the high impact that the isolation method can have on such analyses.

Next, we evaluated whether we could identify genes for which results would differ dramatically between methods. Surprisingly, pathogenic mutations in the CREB-binding protein (CREBBP, also known as CBP) gene were abundantly detected in our dataset. CREBBP is a transcriptional coactivator frequently mutated in hematological malignancies^59, 60^, and inactivating mutations in CREBBP can promote leukemogenesis and predict poor prognosis in AML^59, 61^. While these mutations were detected in all patients, only Ficoll enabled their detection in three patients, and higher variant allele frequencies (VAFs) were observed in Ficoll samples for the other patients (Figure 4I). Because Ficoll increases the proportion of AML blasts, the improved detection of CREBBP mutations possibly reflects the higher fraction of RNA-seq reads originating from leukemic cells.

Another striking observation was the R882 mutation in the DNMT3A gene, which was detected specifically in the Lysis sample of P44 (Figure 4J), in agreement with the reported mutational profile of this patient (Table 1). Manual examination revealed a VAF of 27% with a coverage of 37 reads in the Lysis sample, while VAF was 8% with a coverage of 13 reads in the Ficoll sample. The low VAF and coverage in the Ficoll sample probably prevented the adequate calling of the variant. Interestingly, a recent report showed that R882 mutations promoted a greater differentiation of hematopoietic stem cells into neutrophils^62^. We therefore hypothesize that the R882 mutation detected herein was expressed in neutrophils (in addition to blasts), which enabled a better detection of the mutation in the Lysis sample (this patient presented low blast percentages in both Lysis and Ficoll samples, with 1.71 and 4.85%, respectively). Accordingly, patients with R882 mutations in the BEAT-AML cohort also showed higher levels of circulating neutrophils, as measured by flow cytometry (Figure 4K).

### Limited impact on immunogenomic analyses

The availability of large RNA-seq datasets has spurred studies of immune cell abundance, particularly T cells, in the blood and bone marrow of AML patients^18, 20, 63^. These analyses are typically performed using deconvolution tools such as CIBERSORTx. We examined whether the leukocyte isolation method could influence such analyses. Because immunogenomic studies rely on relative comparisons rather than absolute quantification, any detrimental effect would alter patient rankings by immune cell abundance. We therefore compared CIBERSORTx results from Lysis and Ficoll samples with flow cytometry data from Lysis samples. As shown in Figure 5, both methods showed good ranking concordance for CD8^+^ and CD4^+^ T cells, B cells, monocytes, and granulocytes, with slightly better agreement for Ficoll. In contrast, NK cell rankings were poor with both methods.

**Figure 5.**
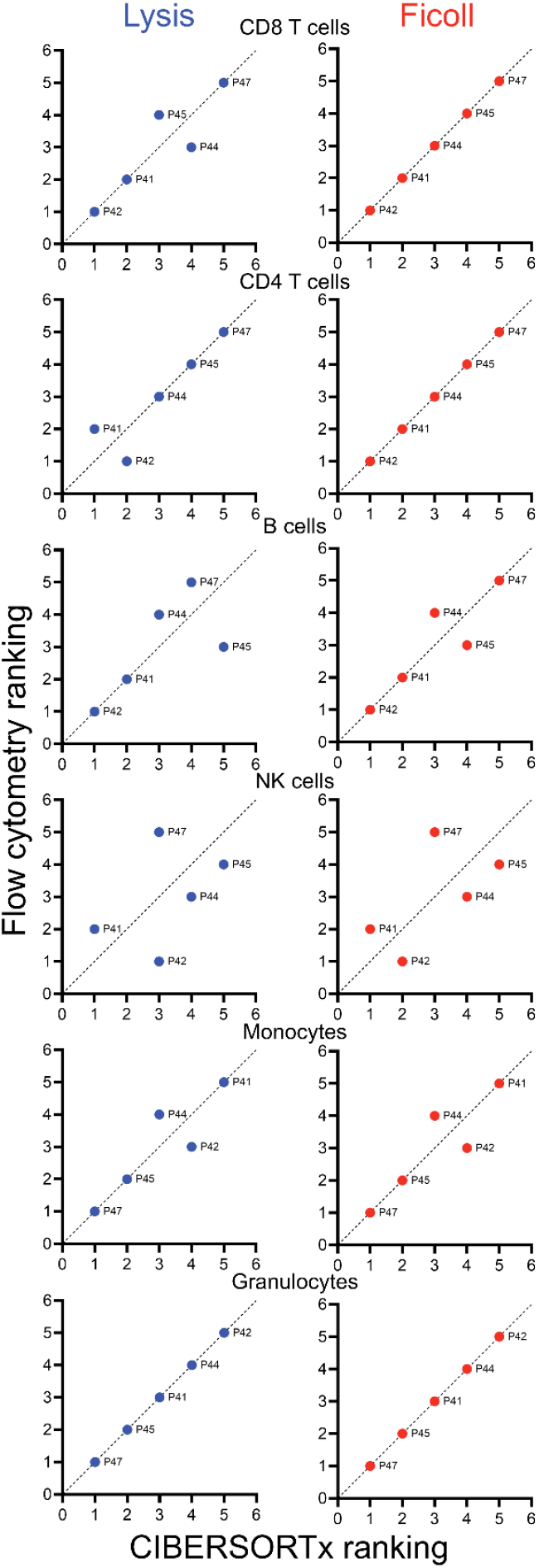
Low impact of the isolation method on immunogenomic analyses. CIBERSORTx deconvolutions performed on RNA-seq data of leukocytes isolated with each method were used to rank samples from patients, and this ranking was compared to the ranking obtained from flow cytometry analyses performed on Lysis samples (the least biased method for measuring the frequency of each cell population). Samples deviating from the identity diagonal line have discordant rankings between RNA-seq and flow cytometry analyses.

## DISCUSSION

Our study demonstrates that Ficoll-based density gradient centrifugation enriches for leukemic blasts and lymphocytes while depleting SSC^high^ granulocytes in AML samples. The distortion introduced by Ficoll isolation had mild to severe impacts on all methodologies tested. We therefore advocate for the systematic use of hemolysis for AML sample processing. In the following sections, we discuss the methodologies explored herein and provide recommendations for each.

Flow cytometry is the most broadly used method to characterize the phenotype of AML blasts in the clinic as well as to monitor MRD. While hemolysis is the standard method recommended by the ELN to process AML samples, Ficoll isolation is still broadly used, especially in research. Here, we observed that Ficoll systematically increased the abundance of SSC^low^ AML blasts making the evaluation of blast abundance largely unreliable. The impact of Ficoll isolation on AML characterization has previously only been explored with respect to blast phenotype^64^, where no differences were detected. However, only binary marker positivity was assessed and not quantitative shifts in population proportions, as performed herein. Importantly, certain AML subtypes, such as KMT2A-rearranged AML and acute promyelocytic leukemia, typically display high SSC values^4^, suggesting that their blast composition (unfortunately not represented in our dataset) may be particularly susceptible to Ficoll-induced distortion. Moreover, chemotherapy has been shown to induce eosinophil-like SSC^high^ AML-derived cells capable of mediating relapse^65^. Therefore, the use of the hemolysis approach is highly recommended for flow cytometry analyses. In our hands, the main limitation to this approach was the abundance of debris. Since debris can be gated out post-acquisition as CD45-negative cells, it does not pose a major issue in flow cytometry analyses.

Regarding xenotransplantations of AML cells, we observed that Ficoll enrichment of T cells led to the development of xenogeneic GVHD symptoms. In contrast, Lysis-isolated samples allowed successful AML engraftment. While we believe that no major differences between isolation methods would have been observed had we used samples containing >80% blasts, our findings highlight the potential risks associated with Ficoll-induced alterations in sample composition. We therefore recommend the systematic use of hemolysis isolation, combined with lymphocyte depletion as previously described^25^, in AML xenograft models to prevent GVHD. Notably, such depletion can also accelerate AML engraftment, even when T cell numbers are below the threshold required to induce GVHD^66^. Importantly, T cell depletion is preferable to positive selection of AML blasts (e.g., CD34^+^ selection) because leukemia-initiating cells can reside in multiple subpopulations, including CD34^−^ cells^23^, and antibody binding to leukemic cells may impair engraftment efficiency in mice^24^.

In contrast with other analytical methods, no striking impact of the isolation methods was observed on *ex vivo* AML cultures. Although Lysis samples showed a slight delay in proliferation compared to Ficoll samples, cellular phenotype and response to chemotherapy were similar. We surmise that this delay results from the lower relative abundance of AML blasts in the Lysis samples. It is also possible that the greater amount of cellular debris present in Lysis samples negatively affects the AML blasts. While this hypothesis cannot be fully excluded, the absence of a viability difference between the two culture conditions suggests that the debris did not induce significant toxicity to the blasts. While it may be tempting to perform a viable-cell sorting step before culture to eliminate debris, this could impose unnecessary stress on the blasts, potentially causing more deleterious than beneficial effects. Finally, the greater size and granularity observed in AML cells surviving chemotherapy (drug-tolerant persisters) suggest that the Ficoll isolation method could affect the composition of this subpopulation. Since drug-tolerant persisters are considered to represent the main population constituting MRD and mediating subsequent relapse, the use of Ficoll in research focused on these cells is not recommended. In summary, we recommend the use of the hemolysis approach to isolate AML cells before *ex vivo* expansion without further fractionation of the samples.

As for other analyses, the main differences in transcriptomic analyses could be attributed to fewer granulocytes and greater lymphocytes and AML blasts by the Ficoll method. The presence of residual red blood cells in Ficoll samples is another possible source of variation. Another striking observation was the non-detection of the DNMT3A^R882^ mutation in the P44 sample isolated with Ficoll. Previous reports showed that neutrophils in AML are dysfunctional and immunosuppressive, thereby increasing the risks of infection and mitigating anti-AML immunity^67^. Moreover, AML blast differentiation is not completely blocked, allowing AML-specific mutations to be detected in more mature populations, such as monocytes and neutrophils^68, 69^. Accordingly, the non-detection of the DNMT3A mutation in the P44 Ficoll sample possibly resulted from the depletion of neutrophils. Thus, the removal of granulocytes by Ficoll isolation represents a significant loss of information for the RNA-seq-based analysis of AML biology.

In addition, and despite the absence of major effects on immunogenomic rankings in our study, the relative enrichment of lymphocytes and depletion of granulocytes introduces an undeniable bias. Therefore, the Lysis method is also preferable for transcriptomic studies, especially for generating large datasets intended for multiple purposes (e.g., immunogenomics, mutation patterns, etc.). Because Lysis samples can contain substantial debris, we recommend sorting viable leukocytes (CD45^+^7-AAD^─^) to avoid RNA contamination from dying cells. This sorting should be followed immediately by RNA extraction to prevent sorting-induced stress responses from altering gene expression profiles. Finally, since we observed a trend toward lower detection of pathogenic mutations in Lysis-isolated samples, possibly due to reduced coverage of blast-specific mutations, we recommend performing high-depth sequencing (>50 million reads) to ensure full coverage of leukemic gene expression and mutation profiles in Lysis-prepared samples.

In conclusion, our work highlights that leukocyte isolation is not a trivial preparatory step but a critical methodological variable that shapes the biological and translational insights derived from AML samples. Our findings advocate for the systematic use of hemolysis as the preferred method for leukocyte isolation in AML research and highlight the necessity to disclose the isolation method used in the preparation of large-scale omics datasets. Hemolysis preserves the entire leukocyte population, maintains relative proportions of subpopulations, and avoids unpredictable biases introduced by Ficoll.

## ACKNOWLEDGMENTS

We thank Sandra Ormenese and Raafat Stephan from the Imaging-Flow Cytometry platform of the GIGA Institute for their assistance. We thank the genomic platform of the GIGA Institute for RNA sequencing. We thank Alexandra Veloso for proofreading the article.

## AUTHOR CONTRIBUTIONS

ADV, FB, and JC collected the samples. BES, AD, CF, JF, MJ, and LCDC performed the experiments. BES, AD, CF, ADV, OK, GC, SC, and GE analyzed and interpreted the data. GE, AG, FB, and JC designed and supervised the study. BES, AD, CF, and GE wrote the article. All authors revised and edited the article.

## CONFLICTS OF INTEREST

The authors declare no conflict of interest.

## DATA AVAILABILITY

Raw RNA-seq data are openly available through GEO (accession number: GSE313042). The remaining datasets generated during and/or analysed during the current study are available from the corresponding author on reasonable request.

## FUNDING

This work was supported by the FNRS Belgium, the Télévie, the Leon Fredericq Foundation, the FRFS-WELBIO strategic research programme, the Fonds Spéciaux de la Recherche (University of Liège), and the ME-TO-YOU Foundation. BES, AD, CF, MJ, OK, LCDC, ADV, and GC are research fellows at the FNRS. LCDC is supported by a grant of the Recherche Scientifique Luxembourg (RSL) non-profit oganization. GE and FB are research associates at the FNRS.

## ETHICS STATEMENT

Peripheral blood samples were obtained after written informed consent (ethics protocol number TJT2225) from the hematology clinical service of the CHU of Liège (University of Liège). All mouse experimental procedures and protocols used in this article were reviewed and approved by the Institutional Animal Care and Use Ethics Committee of the University of Liège, Belgium. The “Guide for the Care and Use of Laboratory Animals”, prepared by the Institute of Laboratory Animal Resources, National Research Council, and published by the National Academy Press, was followed carefully.

## Notes

https://www.ncbi.nlm.nih.gov/geo/query/acc.cgi?acc=GSE313042

### Competing Interest Statement

The authors have declared no competing interest.

