## Supplementary material for "Biases introduced by Ficoll-based isolation in acute myeloid leukemia sample analyses support the use of hemolysis": Table S1

**Supplementary Table 1.** Differential gene expression analysis comparing Ficoll vs Lysis samples. Genes upregulated are overexpressed in Ficoll samples. Only genes significantly differentially expressed (p<0,05) and above or below our log2(FC) threshold (>1 or <-1) are shown. This analysis was performed with the limma-voom packages in paired mode.

| Gene | log2(FC) | Average Expression | p-value |
|---|---|---|---|
| SPTBN4 | 3,402757382 | -2,330945771 | 0,002910435 |
| HLA-DQA2 | 3,207858724 | -2,75073607 | 0,020593309 |
| CTAGE6 | 2,88324162 | -2,674653521 | 0,013862963 |
| ESPNL | 2,622135521 | -1,411132647 | 0,007295828 |
| NME1-NMI | 2,5191616 | -1,003411935 | 0,024975092 |
| CCL3L3 | 2,328227656 | -0,667933377 | 0,005154296 |
| KRT7 | 2,208998294 | -2,004880307 | 0,024440111 |
| MLPH | 2,175874281 | -1,621057275 | 0,015866228 |
| RGS6 | 2,141949843 | -1,337252479 | 0,005369965 |
| SLC17A7 | 2,123114688 | -1,435061927 | 0,019191112 |
| CENPS-C( | 2,056203313 | -0,867119107 | 0,01899794 |
| EFCAB6 | 2,040329786 | -1,271259499 | 0,016567288 |
| PLEKHA6 | 1,969593101 | -1,197605533 | 0,045312275 |
| RGS11 | 1,936902143 | -1,641698697 | 0,039970769 |
| THEM5 | 1,917227902 | -2,017732729 | 0,015479681 |
| DLC1 | 1,907634412 | 0,37038283 | 0,004922447 |
| DLX4 | 1,879577108 | -1,837257251 | 0,038273293 |
| PCDHGA3 | 1,81841722 | -2,461264868 | 0,03194846 |
| PER2 | 1,765090235 | 0,410657146 | 0,013551556 |
| FAM72D | 1,738739027 | 0,666606942 | 0,032236309 |
| ZNF559-ZI | 1,700459256 | 0,290642581 | 0,026578182 |
| C1QTNF3- | 1,697727935 | -0,527331769 | 0,024727474 |
| CCL3 | 1,613034883 | 1,601056196 | 0,016375318 |
| KLC3 | 1,611944292 | 0,374057293 | 0,009514619 |
| NAPSA | 1,586036811 | -0,581396345 | 0,029965021 |
| TMEM181 | 1,529916309 | -0,602958924 | 0,027297365 |
| KRT79 | 1,529323828 | -2,865905086 | 0,043927435 |
| PGM3 | 1,515165537 | 2,017117284 | 0,018796036 |
| CXCR6 | 1,491615772 | -0,115309035 | 0,04987426 |
| AHSP | 1,47735309 | 1,598741647 | 0,000413885 |
| NEIL2 | 1,396681115 | 0,46533066 | 0,043567815 |
| IFIT1B | 1,371652942 | 0,66109679 | 0,002147128 |
| S100B | 1,370803803 | 0,213218713 | 0,004602078 |
| HEYL | 1,361324036 | -0,129956112 | 0,017755916 |
| KRT1 | 1,355087342 | 1,176040674 | 0,000917623 |
| CELF6 | 1,35272488 | 1,553366119 | 0,006396875 |
| PARD6B | 1,334037517 | 0,634451519 | 0,022996586 |
| CACNA2D | 1,328973815 | -1,398440131 | 0,002956243 |
| GYPE | 1,305071149 | -0,154927191 | 0,014313029 |
| RBM11 | 1,300860612 | -1,233410341 | 0,048965715 |
| GPC2 | 1,277236701 | 1,392792523 | 0,006553184 |
| KCNE5 | 1,229398331 | -1,27978675 | 0,036212167 |
| ARHGEF3 | 1,202414862 | -0,70821957 | 0,023271007 |

| | | | |
|---|---|---|---|
| ADAT3 | 1,201688264 | 1,45925494 | 0,008266665 |
| DOC2A | 1,180951038 | -0,303753986 | 0,041075752 |
| HBM | 1,180096806 | 1,599923964 | 0,007710605 |
| TRIM10 | 1,177037108 | 0,066294101 | 0,005483582 |
| HEMGN | 1,16942603 | 3,140116689 | 0,002047561 |
| CA1 | 1,144646543 | 4,244940905 | 0,010505255 |
| ITGAV | 1,140979147 | 2,03854029 | 0,012300124 |
| KLHDC8A | 1,129015711 | -2,502870695 | 0,03463384 |
| A4GALT | 1,112502289 | -1,176486897 | 0,041542816 |
| EDIL3 | 1,108577352 | -1,620718463 | 0,003194099 |
| GYPB | 1,104386777 | -1,367040942 | 0,002005449 |
| SMIM5 | 1,088477219 | 1,228005041 | 0,021014346 |
| MFSD2B | 1,084966611 | 0,962805791 | 0,004398611 |
| TMOD1 | 1,08317041 | 2,285316146 | 0,006202429 |
| SPTB | 1,066689827 | 3,005961542 | 0,000346614 |
| GORASP1 | 1,055387723 | 4,063820169 | 0,017776847 |
| DLX6 | -1,000400778 | 0,440809809 | 0,042511859 |
| AVIL | -1,010546057 | 2,47223861 | 0,004330682 |
| PTP4A3 | -1,014110366 | 5,309903646 | 0,00457792 |
| FOLR3 | -1,014139776 | 2,099078747 | 0,03281322 |
| PNPLA1 | -1,021738261 | 2,400849612 | 0,01037134 |
| REM2 | -1,022237488 | 2,109695054 | 0,00728919 |
| LIPN | -1,023199024 | 2,86742627 | 0,011684996 |
| GRAMD1C | -1,02464506 | 1,390520992 | 0,025650824 |
| CCDC153 | -1,026157105 | 1,120508384 | 0,046204996 |
| TREML2 | -1,031168912 | 4,977338885 | 6,08164E-05 |
| TPST1 | -1,036336762 | 2,303572776 | 0,007056885 |
| RGS2 | -1,039903478 | 7,902841511 | 0,015767747 |
| RYBP | -1,047941153 | 1,99414586 | 0,023432495 |
| THBD | -1,054226922 | 2,954798613 | 0,026340175 |
| IFITM2 | -1,056518499 | 7,424495915 | 2,91845E-05 |
| NEK5 | -1,059125906 | 0,890389765 | 0,030269712 |
| SCN1B | -1,059803204 | 1,45184985 | 0,030844578 |
| CYP3A5 | -1,062609712 | 0,796174106 | 0,027580076 |
| H2BC5 | -1,068515378 | 2,286926184 | 0,02157709 |
| CA4 | -1,077036961 | 1,613663789 | 0,007493826 |
| CSF3R | -1,084254247 | 8,134530109 | 0,015013368 |
| CEP19 | -1,091497177 | 2,784875739 | 0,002527953 |
| FCGR2A | -1,094219627 | 5,923238623 | 0,025740951 |
| TMED7-TI( | -1,096173988 | 2,131204576 | 0,017882237 |
| H2AC6 | -1,097550319 | 5,209533065 | 0,000270289 |
| FPR1 | -1,101798025 | 6,829162034 | 0,021813989 |
| CEACAM3 | -1,103161887 | 2,853137112 | 0,001772907 |
| TMCC3 | -1,109172391 | 3,90264579 | 0,008465066 |
| CCDC96 | -1,111093835 | 1,963615394 | 0,001961508 |
| CLEC4E | -1,120043809 | 3,870882769 | 0,005518214 |
| TNFSF14 | -1,12035571 | 3,672577967 | 0,007346855 |
| CXCL16 | -1,121572848 | 4,025462779 | 0,02430712 |

| | | | |
|---|---|---|---|
| C10orf105 | -1,136192693 | 0,872902821 | 0,029412256 |
| BASP1 | -1,141093168 | 6,840271485 | 0,00849532 |
| VWA7 | -1,146339349 | 0,809940439 | 0,026329325 |
| HSPA6 | -1,162670513 | 4,957801747 | 0,004196724 |
| CHDH | -1,170298519 | 0,892900461 | 0,016223062 |
| ABCA10 | -1,170550977 | -0,006394168 | 0,028424677 |
| CREB5 | -1,171974427 | 4,067030953 | 0,014439457 |
| CCR3 | -1,172809395 | 1,814964211 | 0,016527661 |
| MANSC1 | -1,174691295 | 2,521863624 | 0,003570963 |
| CDC42EP2 | -1,200040972 | 2,262823592 | 0,024487671 |
| TNFRSF12 | -1,201756434 | 0,355177282 | 0,015460552 |
| S100A9 | -1,203631226 | 10,6858931 | 0,038684602 |
| SERPINA1 | -1,205372224 | 7,722174866 | 0,036767476 |
| TRIB3 | -1,210548304 | 1,866407604 | 0,014078637 |
| SLC35F3 | -1,222221528 | -0,917466821 | 0,040498037 |
| IRAG1 | -1,236004345 | 3,124095378 | 0,000880685 |
| TPRG1 | -1,236437435 | 0,026871618 | 0,023235819 |
| VNN2 | -1,245981163 | 6,748448126 | 0,006351413 |
| OR6Y1 | -1,250877014 | 0,067969871 | 0,016787842 |
| CXCL1 | -1,259608347 | 2,822825573 | 0,00822499 |
| SUSD2 | -1,262082208 | 0,120325571 | 0,026294843 |
| ADM | -1,274398752 | 3,25891047 | 0,02289183 |
| CCDC141 | -1,280879035 | 0,158562837 | 0,046977848 |
| PRRT1 | -1,304062639 | -0,518558054 | 0,02155115 |
| DUSP1 | -1,306546842 | 8,689075466 | 0,030786668 |
| TREM1 | -1,308841594 | 5,528958346 | 0,019954753 |
| FOS | -1,3127995 | 9,151444375 | 0,045784605 |
| HCAR2 | -1,319787806 | 1,879656562 | 0,007098296 |
| INKA2 | -1,321601634 | 4,591663597 | 0,000877155 |
| KREMEN1 | -1,332747345 | 3,169184229 | 0,01077926 |
| MTMR7 | -1,333154883 | -1,508367259 | 0,014016262 |
| SPP1 | -1,35315591 | -0,966413897 | 0,031513875 |
| TMPRSS9 | -1,363625106 | 0,10576279 | 0,045426615 |
| TMEM272 | -1,375256176 | 0,979624307 | 0,004586873 |
| DEFA1B | -1,377123297 | -0,678542841 | 0,001254714 |
| KCNJ2 | -1,379213371 | 2,42951648 | 0,01301745 |
| IZUMO1 | -1,413834488 | -1,555925931 | 0,042573641 |
| PLIN5 | -1,427073004 | 2,172025939 | 0,007608897 |
| RUBCNL | -1,441714923 | 3,872033655 | 0,006776973 |
| XKR7 | -1,461836678 | -1,153818017 | 0,014985221 |
| SPESP1 | -1,467540807 | -2,049626204 | 0,04567364 |
| MGAM | -1,490835111 | 6,825098434 | 0,004091787 |
| H4C8 | -1,5000652 | 2,091790897 | 0,006867993 |
| TSPAN6 | -1,506920714 | -0,570628039 | 0,039766517 |
| KNDC1 | -1,511484607 | 1,429976305 | 0,005382736 |
| RIPK4 | -1,519491624 | -1,60186552 | 0,011818385 |
| NUAK1 | -1,520298758 | -1,041102554 | 0,017405261 |
| ADGRE3 | -1,546402845 | 3,574670833 | 0,00021295 |

| | | | |
|---|---|---|---|
| PHOSPHC | -1,547041787 | 3,350696128 | 0,00018459 |
| ARHGAP4 | -1,566756337 | -0,733349467 | 0,041038399 |
| DUSP2 | -1,581212996 | 3,608216929 | 0,00021781 |
| STUM | -1,58188727 | -1,732893488 | 0,007789064 |
| SLC2A10 | -1,593734422 | -1,004653339 | 0,046875427 |
| ZNF608 | -1,62347049 | 2,074397086 | 0,000124644 |
| NPIPB15 | -1,636636967 | -1,296848373 | 0,035074196 |
| H2BC4 | -1,638705736 | 1,585399957 | 0,00272982 |
| SMPD3 | -1,67754465 | 1,720312329 | 0,007594947 |
| NAMPT | -1,681745419 | 8,744303898 | 0,000709527 |
| CCNJL | -1,681948943 | 1,728826628 | 0,000593033 |
| ZFP91-CN | -1,690180224 | 0,957312103 | 0,018034467 |
| KCNMA1 | -1,692794319 | 0,167422234 | 0,010465668 |
| CASP5 | -1,698372326 | 0,936028708 | 0,02966194 |
| CPLX2 | -1,715225642 | -2,197297054 | 0,017274812 |
| PM20D1 | -1,721378981 | -1,607217757 | 0,021164753 |
| AQP9 | -1,726354586 | 5,959742095 | 0,008458645 |
| STEAP4 | -1,732318328 | 4,916516663 | 0,001124118 |
| MMP25 | -1,752362761 | 5,07121608 | 0,015419492 |
| DSC2 | -1,761595792 | -1,077033133 | 0,003638193 |
| PLIN4 | -1,768854082 | 1,817511512 | 0,009351811 |
| ARMH2 | -1,806601096 | -0,735271672 | 0,012397075 |
| BTNL8 | -1,808342097 | 1,732570304 | 0,000272485 |
| RAI2 | -1,809414501 | -1,52982422 | 0,015927247 |
| FFAR4 | -1,820743267 | -1,113729096 | 0,017762391 |
| CMTM2 | -1,841127324 | 2,370024571 | 0,001061421 |
| HBG1 | -1,844029968 | -1,982572456 | 0,005219575 |
| LRRC37A | -1,848658545 | -0,826027026 | 0,035710221 |
| CLDN10 | -1,849225338 | 1,692682063 | 0,009099149 |
| TNFRSF1( | -1,865008599 | 4,594403725 | 0,00021778 |
| PNMA6A | -1,870149147 | -0,581703888 | 0,015422783 |
| CACNG5 | -1,873300293 | -1,714903673 | 0,01372591 |
| KCNJ15 | -1,886367603 | 3,594809674 | 0,000263265 |
| NINL | -1,898361023 | -1,570480201 | 0,005256738 |
| CCR4 | -1,899232834 | -1,051730391 | 0,035882456 |
| GBP6 | -1,903614883 | -1,127575371 | 0,021237082 |
| FFAR2 | -1,920127138 | 3,91014501 | 0,00355714 |
| MME | -1,92106149 | 4,76662646 | 0,000196244 |
| AOC3 | -1,936182568 | 0,480919491 | 0,002736846 |
| CXCR2 | -1,936663308 | 5,610758985 | 0,000127068 |
| SLCO1A2 | -1,954289159 | -1,208992846 | 0,039537522 |
| KSR2 | -1,960482712 | -1,644498878 | 0,024850131 |
| MAK | -1,991731526 | 0,155774117 | 0,025388165 |
| ITPRID1 | -1,99543248 | -1,455617297 | 0,035070125 |
| HAMP | -1,998268489 | -0,498147668 | 0,004157711 |
| CFAP53 | -2,040415018 | -0,786193716 | 0,021252503 |
| TGM3 | -2,077677731 | 0,393009878 | 0,000764903 |
| KRT2 | -2,116424289 | -1,770867038 | 0,019062211 |

| | | | |
|---|---|---|---|
| SLC22A1 | -2,129515685 | -0,629022264 | 0,029138421 |
| IL1R2 | -2,140045378 | 4,077994068 | 0,000704398 |
| TVP23C-C | -2,157083935 | 0,014916781 | 0,026312601 |
| H2BC15 | -2,176515223 | 0,057847566 | 0,00115504 |
| GPR82 | -2,184425935 | -1,719739346 | 0,005957681 |
| PIP5KL1 | -2,192387129 | -0,658600309 | 0,010862754 |
| KIAA0319 | -2,228896272 | -0,980208557 | 0,021296646 |
| CD177 | -2,23824401 | 2,042368091 | 0,027277892 |
| CXCR1 | -2,244678011 | 4,442402931 | 2,9255E-05 |
| FCGR3B | -2,256362942 | 6,105889764 | 0,000151712 |
| KRT23 | -2,258491333 | 1,42523469 | 1,84836E-05 |
| AOC2 | -2,270093565 | 0,823764648 | 0,00719394 |
| RNF103-C | -2,291674431 | -1,299758922 | 0,016243416 |
| SH3BGR | -2,359379294 | -0,915573858 | 0,003230248 |
| ALOX15 | -2,48986205 | -0,144996603 | 0,00026322 |
| PRSS33 | -2,522442524 | -1,743026309 | 0,031777083 |
| LSMEM1 | -2,523740194 | -0,992609889 | 0,045611755 |
| MYBPH | -2,53395824 | -2,194480535 | 0,001975352 |
| LPIN3 | -2,59666692 | -2,34569281 | 0,013295385 |
| PI3 | -2,762982174 | 2,595140112 | 0,000256306 |
| NECAB2 | -3,111893701 | -1,737483161 | 0,002584952 |
| HEY1 | -3,182889912 | -0,616739209 | 1,29644E-06 |
| ALPL | -3,315805126 | 3,6309588 | 9,68903E-06 |
| TSPEAR | -3,492207396 | -2,199878491 | 0,000169552 |
