## Supplementary material for "Biases introduced by Ficoll-based isolation in acute myeloid leukemia sample analyses support the use of hemolysis": Table S2

| **Supplementary Table 2.** Details of mutations detected by Freebayes and annotated as pathogenic or likely pathogenic in ClinVar. | | | | | | | | | |
|---|---|---|---|---|---|---|---|---|---|
| Patient | Condition | Chromosome | Position | Reference | Alternative | #alt reads | Coverage | VAF | Gene |
| P41 | Ficoll | chr3 | 33014057 | T | C | 54 | 121 | 44,63 | GLB1 |
| P41 | Ficoll | chr3 | 43717675 | G | A | 6 | 68 | 8,82 | ABHD5 |
| P41 | Ficoll | chr3 | 49122045 | G | A | 5 | 16 | 31,25 | LAMB2 |
| P41 | Ficoll | chr3 | 75741671 | A | G | 7 | 7 | 100 | ZNF717 |
| P41 | Ficoll | chr6 | 166948626 | C | A | 19 | 33 | 57,58 | RNASET2 |
| P41 | Ficoll | chr9 | 22003368 | G | A | 6 | 6 | 100 | CDKN2B |
| P41 | Ficoll | chr11 | 126275413 | C | T | 13 | 39 | 33,33 | FOXRED1 |
| P41 | Ficoll | chr12 | 25245350 | C | A | 6 | 31 | 19,35 | KRAS |
| P41 | Ficoll | chr14 | 87992996 | C | G | 5 | 83 | 6,02 | GALC |
| P41 | Ficoll | chr16 | 3729448 | G | A | 7 | 62 | 11,29 | CREBBP |
| P41 | Ficoll | chr16 | 3736739 | G | T | 9 | 50 | 18 | CREBBP |
| P41 | Ficoll | chr16 | 3793494 | G | A | 10 | 33 | 30,3 | CREBBP |
| P41 | Ficoll | chr16 | 89761995 | C | T | 16 | 24 | 66,67 | FANCA |
| P41 | Ficoll | chr17 | 80108716 | T | C | 8 | 108 | 7,41 | GAA |
| P41 | Ficoll | chr17 | 80369794 | T | C | 6 | 38 | 15,79 | RNF213 |
| P41 | Ficoll | chr19 | 33387892 | C | T | 12 | 53 | 22,64 | PEPD |
| P41 | Lysis | chr1 | 19227402 | G | A | 6 | 49 | 12,24 | EMC1 |
| P41 | Lysis | chr3 | 33014057 | T | C | 82 | 133 | 61,65 | GLB1 |
| P41 | Lysis | chr3 | 167684361 | G | A | 6 | 63 | 9,52 | PDCD10 |
| P41 | Lysis | chr6 | 166948626 | C | A | 8 | 16 | 50 | RNASET2 |
| P41 | Lysis | chr8 | 18062422 | A | G | 6 | 61 | 9,84 | ASAH1 |
| P41 | Lysis | chr9 | 22003368 | G | A | 22 | 22 | 100 | CDKN2B |
| P41 | Lysis | chr11 | 64804748 | CTC | CC | 6 | 34 | 17,65 | MEN1 |
| P41 | Lysis | chr12 | 25245350 | C | A | 8 | 40 | 20 | KRAS |
| P41 | Lysis | chr17 | 16006475 | C | T | 14 | 33 | 42,42 | TTC19 |
| P41 | Lysis | chr19 | 11033384 | T | C | 6 | 59 | 10,17 | SMARCA4 |
| P41 | Lysis | chr19 | 11105588 | G | A | 6 | 51 | 11,76 | LDLR |
| P41 | Lysis | chr20 | 35508113 | C | T | 6 | 26 | 23,08 | CEP250 |
| P41 | Lysis | chrX | 19355385 | T | C | 6 | 97 | 6,19 | PDHA1 |
| P41 | Lysis | chrX | 154349360 | A | T | 8 | 9 | 88,89 | FLNA |
| P42 | Ficoll | chr1 | 23693914 | G | A | 5 | 19 | 26,32 | RPL11 |
| P42 | Ficoll | chr1 | 114716126 | C | T | 33 | 63 | 52,38 | NRAS |
| P42 | Ficoll | chr1 | 156134911 | G | A | 8 | 74 | 10,81 | LMNA |
| P42 | Ficoll | chr2 | 47478303 | G | T | 5 | 35 | 14,29 | MSH2 |
| P42 | Ficoll | chr3 | 128481275 | C | T | 258 | 397 | 64,99 | GATA2 |
| P42 | Ficoll | chr6 | 166948626 | C | A | 7 | 10 | 70 | RNASET2 |
| P42 | Ficoll | chr10 | 89225168 | A | G | 6 | 82 | 7,32 | LIPA |
| P42 | Ficoll | chr11 | 5226928 | ACC | CCT,GCC | 126,75 | 1326 | 9,56 | HBB |
| P42 | Ficoll | chr11 | 62150080 | G | C | 39 | 69 | 56,52 | INCENP |
| P42 | Ficoll | chr12 | 31089147 | G | A | 16 | 85 | 18,82 | DDX11 |
| P42 | Ficoll | chr16 | 3736739 | G | T | 8 | 69 | 11,59 | CREBBP |
| P42 | Ficoll | chr16 | 28491758 | A | G | 10 | 61 | 16,39 | CLN3 |
| P42 | Ficoll | chr17 | 80117702 | G | A | 5 | 27 | 18,52 | GAA |
| P42 | Ficoll | chr20 | 10644356 | C | A | 6 | 6 | 100 | JAG1 |
| P42 | Ficoll | chr22 | 20994271 | T | G | 7 | 11 | 63,64 | LZTR1 |
| P42 | Ficoll | chrX | 41216199 | G | A | 7 | 110 | 6,36 | USP9X |
| P42 | Lysis | chr1 | 114716126 | C | T | 28 | 53 | 52,83 | NRAS |
| P42 | Lysis | chr1 | 156593885 | G | A | 6 | 39 | 15,38 | NAXE |
| P42 | Lysis | chr3 | 36993623 | C | T | 18 | 71 | 25,35 | MLH1 |
| P42 | Lysis | chr3 | 128481275 | C | T | 292 | 368 | 79,35 | GATA2 |
| P42 | Lysis | chr3 | 128486072 | T | G | 6 | 98 | 6,12 | GATA2 |
| P42 | Lysis | chr6 | 166948626 | C | A | 11 | 26 | 42,31 | RNASET2 |
| P42 | Lysis | chr7 | 157367402 | T | A | 11 | 170 | 6,47 | DNAJB6 |
| P42 | Lysis | chr8 | 42288691 | C | T | 6 | 67 | 8,96 | IKBKB |
| P42 | Lysis | chr12 | 31089147 | G | A | 9 | 68 | 13,24 | DDX11 |
| P42 | Lysis | chr12 | 53308095 | G | A | 6 | 70 | 8,57 | AAAS |
| P42 | Lysis | chr12 | 115987147 | C | T | 5 | 89 | 5,62 | MED13L |

| | | | | | | | | | |
|---|---|---|---|---|---|---|---|---|---|
| P42 | Lysis | chr14 | 94382590 | A | G | 7 | 44 | 15,91 | SERPINA1 |
| P42 | Lysis | chr21 | 46116028 | G | A | 6 | 31 | 19,35 | COL6A2 |
| P44 | Ficoll | chr1 | 23693914 | G | A | 6 | 22 | 27,27 | RPL11 |
| P44 | Ficoll | chr1 | 26767868 | T | G | 5 | 65 | 7,69 | ARID1A |
| P44 | Ficoll | chr1 | 35893730 | T | C | 6 | 16 | 37,5 | AGO1 |
| P44 | Ficoll | chr1 | 215586516 | G | A | 7 | 7 | 100 | KCTD3 |
| P44 | Ficoll | chr3 | 136327184 | C | T | 6 | 24 | 25 | PCCB |
| P44 | Ficoll | chr5 | 177280700 | T | C | 5 | 73 | 6,85 | NSD1 |
| P44 | Ficoll | chr6 | 166948626 | C | A | 29 | 47 | 61,7 | RNASET2 |
| P44 | Ficoll | chr11 | 2571328 | T | C | 9 | 54 | 16,67 | KCNQ1 |
| P44 | Ficoll | chr11 | 5226928 | ACC | CCT,GCC | 6,5 | 11 | 59,09 | HBB |
| P44 | Ficoll | chr11 | 108316041 | G | A | 16 | 60 | 26,67 | ATM |
| P44 | Ficoll | chr12 | 6587885 | C | T | 5 | 40 | 12,5 | CHD4 |
| P44 | Ficoll | chr12 | 31089147 | G | A | 5 | 62 | 8,06 | DDX11 |
| P44 | Ficoll | chr14 | 94380925 | T | A | 707 | 1321 | 53,52 | SERPINA1 |
| P44 | Ficoll | chr14 | 95096633 | C | T | 12 | 96 | 12,5 | DICER1 |
| P44 | Ficoll | chr15 | 40407639 | C | T | 5 | 74 | 6,76 | IVD |
| P44 | Ficoll | chr16 | 2087918 | T | C | 10 | 36 | 27,78 | TSC2 |
| P44 | Ficoll | chr16 | 3736739 | G | T | 10 | 50 | 20 | CREBBP |
| P44 | Ficoll | chr16 | 89280568 | T | G | 15 | 28 | 53,57 | ANKRD11 |
| P44 | Ficoll | chr17 | 75758286 | G | A | 5 | 62 | 8,06 | GALK1 |
| P44 | Ficoll | chr22 | 20994271 | T | C | 6 | 20 | 30 | LZTR1 |
| P44 | Ficoll | chrX | 101353268 | G | A | 10 | 95 | 10,53 | BTK |
| P44 | Ficoll | chrX | 101356910 | A | G | 7 | 110 | 6,36 | BTK |
| P44 | Lysis | chr1 | 23693914 | G | A | 8 | 30 | 26,67 | RPL11 |
| P44 | Lysis | chr1 | 226982734 | G | A | 9 | 109 | 8,26 | COQ8A |
| P44 | Lysis | chr1 | 235693370 | G | A | 9 | 45 | 20 | LYST |
| P44 | Lysis | chr2 | 25234373 | C | T | 10 | 37 | 27,03 | DNMT3A |
| P44 | Lysis | chr2 | 47801003 | G | A | 10 | 36 | 27,78 | MSH6 |
| P44 | Lysis | chr2 | 74462273 | G | A | 10 | 74 | 13,51 | MOGS |
| P44 | Lysis | chr2 | 165911404 | A | G | 5 | 18 | 27,78 | TTC21B |
| P44 | Lysis | chr3 | 134559198 | C | A | 6 | 6 | 100 | CEP63 |
| P44 | Lysis | chr6 | 137206214 | A | G | 5 | 74 | 6,76 | IFNGR1 |
| P44 | Lysis | chr6 | 166948626 | C | A | 16 | 26 | 61,54 | RNASET2 |
| P44 | Lysis | chr10 | 88947370 | A | G | 10 | 35 | 28,57 | ACTA2 |
| P44 | Lysis | chr12 | 31089147 | G | A | 9 | 33 | 27,27 | DDX11 |
| P44 | Lysis | chr14 | 94380925 | T | A | 669 | 1131 | 59,15 | SERPINA1 |
| P44 | Lysis | chr16 | 3254736 | C | T | 6 | 75 | 8 | MEFV |
| P44 | Lysis | chr16 | 3729409 | G | A | 6 | 88 | 6,82 | CREBBP |
| P44 | Lysis | chr16 | 3736739 | G | T | 8 | 91 | 8,79 | CREBBP |
| P44 | Lysis | chr16 | 20737873 | G | A | 6 | 45 | 13,33 | ACSM3 |
| P44 | Lysis | chr16 | 89280568 | T | G | 19 | 28 | 67,86 | ANKRD11 |
| P44 | Lysis | chr16 | 89739476 | A | G | 6 | 108 | 5,56 | FANCA |
| P44 | Lysis | chr17 | 5583728 | A | G | 8 | 78 | 10,26 | NLRP1 |
| P44 | Lysis | chr19 | 1401317 | C | G | 5 | 7 | 71,43 | GAMT |
| P44 | Lysis | chr22 | 20988032 | T | G | 8 | 28 | 28,57 | LZTR1 |
| P44 | Lysis | chrX | 53399637 | C | T | 7 | 110 | 6,36 | SMC1A |
| P44 | Lysis | chrX | 153729266 | A | G | 5 | 61 | 8,2 | ABCD1 |
| P45 | Ficoll | chr1 | 17027787 | G | A | 6 | 118 | 5,08 | SDHB |
| P45 | Ficoll | chr2 | 27312499 | G | A | 5 | 89 | 5,62 | MPV17 |
| P45 | Ficoll | chr2 | 74092933 | T | C | 8 | 13 | 61,54 | TET3 |
| P45 | Ficoll | chr3 | 10142040 | T | C | 12 | 217 | 5,53 | VHL |
| P45 | Ficoll | chr3 | 75741671 | A | G | 10 | 14 | 71,43 | ZNF717 |
| P45 | Ficoll | chr3 | 179234302 | G | T | 7 | 43 | 16,28 | PIK3CA |
| P45 | Ficoll | chr5 | 112842417 | A | T | 10 | 26 | 38,46 | APC |
| P45 | Ficoll | chr6 | 31592893 | C | T | 5 | 18 | 27,78 | NCR3 |
| P45 | Ficoll | chr6 | 166948626 | C | A | 6 | 10 | 60 | RNASET2 |
| P45 | Ficoll | chr7 | 141641362 | C | T | 6 | 46 | 13,04 | AGK |
| P45 | Ficoll | chr8 | 38424684 | C | T | 7 | 28 | 25 | FGFR1 |
| P45 | Ficoll | chr10 | 111079821 | A | G | 8 | 8 | 100 | ADRA2A |

| | | | | | | | | | |
|---|---|---|---|---|---|---|---|---|---|
| P45 | Ficoll | chr11 | 77379252 | A | G | 6 | 14 | 42,86 | PAK1 |
| P45 | Ficoll | chr12 | 31089147 | G | A | 12 | 43 | 27,91 | DDX11 |
| P45 | Ficoll | chr12 | 65171119 | G | A | 5 | 22 | 22,73 | LEMD3 |
| P45 | Ficoll | chr12 | 112472971 | C | T | 7 | 71 | 9,86 | PTPN11 |
| P45 | Ficoll | chr14 | 20452057 | A | G | 27 | 54 | 50 | OSGEP |
| P45 | Ficoll | chr16 | 3736739 | G | T | 6 | 58 | 10,34 | CREBBP |
| P45 | Ficoll | chr17 | 80215086 | A | G | 6 | 33 | 18,18 | SGSH |
| P45 | Ficoll | chr19 | 36067859 | C | T | 6 | 8 | 75 | WDR62 |
| P45 | Ficoll | chr19 | 49595776 | C | T | 8 | 43 | 18,6 | PRR12 |
| P45 | Ficoll | chrX | 149498153 | A | G | 8 | 156 | 5,13 | IDS |
| P45 | Lysis | chr1 | 11999058 | G | A | 10 | 55 | 18,18 | MFN2 |
| P45 | Lysis | chr1 | 145927328 | C | G | 6 | 7 | 85,71 | RBM8A |
| P45 | Lysis | chr2 | 47799565 | G | T | 5 | 70 | 7,14 | MSH6 |
| P45 | Lysis | chr2 | 88691951 | G | A | 10 | 40 | 25 | RPIA |
| P45 | Lysis | chr2 | 229799343 | G | A | 6 | 49 | 12,24 | TRIP12 |
| P45 | Lysis | chr3 | 158690234 | C | T | 8 | 39 | 20,51 | GFM1 |
| P45 | Lysis | chr4 | 158706317 | G | A | 5 | 32 | 15,63 | ETFDH |
| P45 | Lysis | chr6 | 31668655 | G | T | 5 | 12 | 41,67 | CSNK2B |
| P45 | Lysis | chr7 | 128398529 | A | G | 6 | 69 | 8,7 | IMPDH1 |
| P45 | Lysis | chr10 | 111079821 | A | G | 9 | 9 | 100 | ADRA2A |
| P45 | Lysis | chr11 | 5226928 | A | G | 5 | 7 | 71,43 | HBB |
| P45 | Lysis | chr12 | 31089147 | G | A | 13 | 52 | 25 | DDX11 |
| P45 | Lysis | chr14 | 20452057 | A | G | 38 | 63 | 60,32 | OSGEP |
| P45 | Lysis | chr14 | 102033157 | C | T | 6 | 106 | 5,66 | DYNC1H1 |
| P45 | Lysis | chr16 | 5079078 | T | C | 5 | 19 | 26,32 | ALG1 |
| P45 | Lysis | chr17 | 2676571 | T | A | 8 | 103 | 7,77 | PAFAH1B1 |
| P45 | Lysis | chr17 | 80108512 | T | C | 5 | 37 | 13,51 | GAA |
| P45 | Lysis | chr19 | 18080847 | G | A | 6 | 76 | 7,89 | IL12RB1 |
| P47 | Ficoll | chr1 | 236863553 | C | T | 6 | 25 | 24 | MTR |
| P47 | Ficoll | chr1 | 241504206 | A | G | 8 | 37 | 21,62 | FH |
| P47 | Ficoll | chr3 | 33016800 | A | G | 10 | 38 | 26,32 | GLB1 |
| P47 | Ficoll | chr3 | 37048952 | T | C | 7 | 60 | 11,67 | MLH1 |
| P47 | Ficoll | chr5 | 75416973 | G | A | 14 | 62 | 22,58 | CERT1 |
| P47 | Ficoll | chr6 | 166948626 | C | A | 12 | 17 | 70,59 | RNASET2 |
| P47 | Ficoll | chr7 | 45068469 | C | T | 8 | 96 | 8,33 | CCM2 |
| P47 | Ficoll | chr9 | 69237907 | G | A | 5 | 25 | 20 | TJP2 |
| P47 | Ficoll | chr10 | 69359951 | G | A | 10 | 68 | 14,71 | HK1 |
| P47 | Ficoll | chr10 | 89007762 | G | T | 6 | 39 | 15,38 | FAS |
| P47 | Ficoll | chr11 | 71438986 | G | A | 8 | 33 | 24,24 | DHCR7 |
| P47 | Ficoll | chr14 | 87968380 | C | T | 8 | 14 | 57,14 | GALC |
| P47 | Ficoll | chr14 | 102032419 | G | A | 5 | 88 | 5,68 | DYNC1H1 |
| P47 | Ficoll | chr15 | 72350587 | C | T | 11 | 115 | 9,57 | HEXA |
| P47 | Ficoll | chr15 | 89318623 | G | A | 6 | 66 | 9,09 | POLG |
| P47 | Ficoll | chr16 | 173008 | G | A | 10 | 16 | 62,5 | HBA2 |
| P47 | Ficoll | chr16 | 3736739 | G | T | 9 | 66 | 13,64 | CREBBP |
| P47 | Ficoll | chr16 | 10907209 | C | T | 10 | 29 | 34,48 | CIITA |
| P47 | Ficoll | chr21 | 44333087 | A | G | 6 | 99 | 6,06 | CFAP410 |
| P47 | Ficoll | chr22 | 20991684 | G | A | 5 | 35 | 14,29 | LZTR1 |
| P47 | Ficoll | chr22 | 50529225 | G | A | 8 | 16 | 50 | TYMP |
| P47 | Ficoll | chrX | 154533589 | A | G | 7 | 57 | 12,28 | G6PD |
| P47 | Lysis | chr1 | 40089456 | G | A | 12 | 107 | 11,21 | PPT1 |
| P47 | Lysis | chr2 | 39058765 | A | G | 5 | 46 | 10,87 | SOS1 |
| P47 | Lysis | chr3 | 4814521 | G | A | 8 | 33 | 24,24 | ITPR1 |
| P47 | Lysis | chr5 | 7878116 | G | T | 7 | 26 | 26,92 | MTRR |
| P47 | Lysis | chr6 | 166948626 | C | A | 13 | 18 | 72,22 | RNASET2 |
| P47 | Lysis | chr7 | 30601179 | T | C | 8 | 81 | 9,88 | GARS1 |
| P47 | Lysis | chr7 | 66994286 | TA | AG | 6 | 112 | 5,36 | SBDS |
| P47 | Lysis | chr8 | 24953770 | G | A | 7 | 15 | 46,67 | NEFL |
| P47 | Lysis | chr10 | 825204 | C | A | 5 | 27 | 18,52 | LARP4B |
| P47 | Lysis | chr10 | 78022270 | G | A | 18 | 38 | 47,37 | POLR3A |

| | | | | | | | | | |
|---|---|---|---|---|---|---|---|---|---|
| P47 | Lysis | chr11 | 5226928 | A | G | 20 | 22 | 90,91 | HBB |
| P47 | Lysis | chr11 | 103253341 | C | T | 8 | 8 | 100 | DYNC2H1 |
| P47 | Lysis | chr15 | 80180191 | G | A | 12 | 24 | 50 | FAH |
| P47 | Lysis | chr16 | 1447038 | G | A | 5 | 48 | 10,42 | CLCN7 |
| P47 | Lysis | chr16 | 3736739 | G | T | 9 | 95 | 9,47 | CREBBP |
| P47 | Lysis | chr17 | 2666076 | A | T | 8 | 62 | 12,9 | PAFAH1B1 |
| P47 | Lysis | chr17 | 7675115 | G | T | 7 | 57 | 12,28 | TP53 |
| P47 | Lysis | chr17 | 78126358 | G | A | 12 | 199 | 6,03 | TMC6 |
| P47 | Lysis | chr19 | 52212730 | G | C | 10 | 95 | 10,53 | PPP2R1A |
| P47 | Lysis | chr22 | 41117690 | C | T | 7 | 127 | 5,51 | EP300 |
| P47 | Lysis | chr22 | 50457113 | T | C | 6 | 12 | 50 | SBF1 |
| P47 | Lysis | chrX | 154534345 | C | A | 6 | 57 | 10,53 | G6PD |
